# Characterizing the fragmentation of AlphaFold predictions

**DOI:** 10.64898/2025.12.19.695436

**Authors:** Frédéric Cazals, Edoardo Sarti

## Abstract

The Nobel prize winning program `AlphaFold2` computes plausible structures of (well) folded proteins. The main quality assessment is based on the *predicted Local Distance Difference Test* (pLDDT), a per amino acid confidence score. To enhance quality assessment, we provide novel quantitative measures to identify *coherent* amino acid (a.a.) stretches along the sequence in terms of pLDDT values. These constructions, grounded in standard techniques from topological data analysis and combinatorics, provide a canonical framework for identifying regions along the protein backbone and analyzing their properties, such as their propensity for disorder and their consistency with a null model. The outcome of our analysis can readily be used to select reliable regions/domains within proteins whose pLDDT values span the entire pLDDT range.

## 1 Introduction

### 1.1 AlphaFold2

`AlphaFold2` **and** `AlphaFold-DB`. `AlphaFold2` ‘s ability to reliably predict protein structures from amino acid sequences has ushered in a new era in structural biology [1]. In recognition of this achievement, its leaders D. Hassabis and J. Jumper were jointly awarded the 2024 Nobel Prize in Chemistry. Each `AlphaFold2` prediction is provided as a PDB file, with individual amino acids assigned a confidence score known as pLDDT (predicted Local Distance Difference Test), related to the Local Distance Difference Test (LDDT) [2]. In a reference structure, define a *native contact* as a pair of atoms within distance *r* – called the inclusion radius. The LDDT statistics at threshold *δ* is fraction of native contacts in a prediction conserved within threshold *δ*. The LDDT score is the average of the previous over several values – typically *δ* ∈ {0.5, 1, 2, 4} Å.

During the learning phase, `AlphaFold2` trains a specific head taking as input the *c*_*s*_(= 384) dimensional embeddings of the residues to regress the LDDT values computed w.r.t. crystal structures [1, SI, Algorithm 29]. At inference time, this regressor is reused to estimate the LDDT of the current prediction.) `AlphaFold2` team defined four confidence categories based on pLDDT values: very high (0.9 ≤pLDDT, dark blue), high (0.7≤ pLDDT *<* 0.9, cyan), low (0.5 ≤pLDDT *<* 0.7, yellow), and very low (pLDDT *<* 0.5, orange).

On the experimental side, the Protein Data Bank currently holds approximately 230*K* experimentally determined structures, in contrast to the *∼* 200*M* protein sequences recorded in `UniProtKB`. This stark imbalance highlights the critical role of structure prediction in enabling genome-wide proteomic analysis (SWISS-MODEL Repository [3]). To address this, `AlphaFold2` has been applied at scale, with predictions now available in the AlphaFold Protein Structure Database database–`AlphaFold-DB`, [4] and https://alphafold.ebi.ac.uk. A comprehensive analysis of this database reveals that for many model organisms, 30% to 40% of amino acids in predicted structures fall within the low or very low pLDDT confidence categories (Table S1).

### Existing analysis for AlphaFold2 predictions

From the structural standpoint, at the whole genome level, an embedding of `AlphaFold2` predictions based on shape-mers has been obtained using dimensionality reduction (t-SNE, [5, Fig. 2]). The structural features within `AlphaFold2` predictions have been examined using the `Geometricus` algorithm and associated embeddings[6]. By clustering these embeddings – using non-negative matrix factorization – across 21 predicted proteomes from `AlphaFold-DB`, 250 structural clusters were identified, including 20 major clusters or superfamilies [5]. An enhanced unsupervised clustering algorithm designed aiming at reliably identifying structural domains has also been developed [7], resulting in meta-clusters consistent with clans in `Pfam`.

The pLDDT metric has been thoroughly analyzed, revealing its effectiveness as robustness indicator. On the one hand, it has been observed that the low-pLDDT regions in `AlphaFold2` predictions correlate with observed or predicted intrinsically disordered regions (IDRs) [8]. On the other hand, it has been noticed that high mean pLDDT values are associated with minimal structural differences between `AlphaFold2` and `trRosetta` models [5]. Importantly, using B-factor values, it has been pointed out that pLDDT does not appear to correlate with flexibility [9].

Finally, it should be noted that structure prediction using `AlphaFold2` classically delivers one structure, while varying experimental conditions in structure determination typically yields different structures. It has been argued that for selected cases (typically well folded structures), predictions fall within the set of experimental outcomes, which is satisfactory [10]. The default usage of `AlphaFold2` consists of generating a single structure, which is not compatible with multistate conformations – as illustrated *e*.*g*. with two-state switching proteins [10, 11]. The method capability has been extended by manipulating the multiple sequence alignment (MSA) used, including clustering of sequences [12] and in silico mutations [12]. These methods naturally expand `AlphaFold2`’s scope. Yet, analyzing all outputs by providing further insights into the pLDDT structural heterogeneity remains of interest.

### 1.2 Contribution and paper overview

As just noticed, the precise relationship between pLDDT values and biophysical properties have remained elusive. While the structural heterogeneity in terms of pLDDT values is believed to be important for quality assessment, no quantitative assessment of this heterogeneity / coherence / fragmentation has been proposed. We fill this gap by analyzing the coherence of pLDDT values along the primary sequence. In other words, we address the question of segmenting a protein into coherent stretches of a.a. with *coherent* pLDDT values. Our analysis relies on the stability of connected components of amino acids (a.a.) along the protein sequence, upon building a suitable filtration which consists of inserting the a.a. by decreasing pLDDT values. The results obtained are confronted against a null model analyzed using methods from combinatorics.

Section 2 introduces our statistics, and Section 3 proceeds with results on `AlphaFold-DB`.

## 2 Methods: persistence based analysis and null model

### 2.1 Persistence based analysis on the primary structure

#### Filtrations on a path graph

We aim to evaluate the *coherence*, with respect to a parameter generically denoted as *u*, of the *n* amino acids of a protein sequence in an `AlphaFold2` prediction. Practically, *u* will be the pLDDT, although other choices are possible, like atomic packing measures for example – see the companion paper [13].

We consider the polypeptide chain as a path graph with *n* vertices and *n −*1 edges. From a topological data analysis perspective [14], we use a filtration 𝒢 _*u*_ encoding a sequence of nested subgraphs of this path graph. Here, the calligraphic letter 𝒢 indicates that the filtration is applied to the path graph, and *u* is the value carried by each each amino acid. Upon inserting a.a. by increasing *u* value, we connect the a.a. being processed to its neighbors along the sequence if already inserted–Algorithm S1. The filtration starts with the empty graph and ends with the complete path graph.

As *u* varies, we track the number of connected components (c.c.) in the graph𝒢 _*u*_ using the Union-Find algorithm [15]. This results in a function *v* = *N*_cc_(*u*) representing the number of connected components at each value of *u*, with *ν* in the interval [1, *n*⌈*/*2⌉]. We note in passing that there are 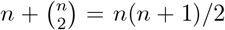 possible connected subgraphs – the *n* singletons plus one connected subgraph for any pair of vertices.

The *persistence* of a connected component in𝒢 _*u*_ is the time elapsed between its birth (value of *u* of the first a.a. inserted) and the moment when it merges with a c.c. that was created earlier. The *persistence diagram* (PD) is the diagram whose *x*-axis (resp. y-axis) is the birth (resp. death) date [14]. The points of the PD are denoted *P* = *{c*_1_, …, *c*_*m*_*}*, and referred to as *critical points* – to be consistent with Morse theory where critical points correspond to topological changes. The persistence of point *c*_*i*_ reads as `p`_𝒢_ (*c*_*i*_) = death_𝒢_ (*c*_*i*_) birth_𝒢_ (*c*_*i*_). We also denote *m*^*′*^ the number of critical points whose persistence is positive–*i*.*e*. critical points away from the diagonal, and define the fraction of positive critical points as 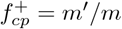

Assuming that the persistences belong to a discrete set 𝒫, we denote ℙ [*·*] the probability mass function *ℙ* : *𝒫 →* [0, 1]. To assess the diversity of persistence values, we consider the normalized entropy of this probability distribution.

Finally, we consider the *salient / persistent* local maxima of the function *N*_cc_(*u*). We identify such maxima using persistence again on the filtration *ℋ*_*v*_ associated with super-level sets 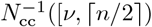 of the function *N*_cc_(*u*). (The calligraphic letter *ℋ* indicates that the filtration is defined on the height function *N*_cc_(*u*).) The persistence `p`_ℋ_ (*c*_*i*_) = death_*ℋ*_ (*c*_*i*_) birth_*ℋ*_ (*c*_*i*_) of a local maximum of *N*_cc_(*u*) is the elevation drop leading to a saddle point also connected to a more elevated local maximum. We simplify the function *N*_cc_(*u*) using the persistence simplification based on the Morse-Smale-Witten chain complex [16]. In particular, we denote PLM(*t*_*v*_) the number of persistent local maxima of *N*_cc_(*u*) at persistence threshold *t*_*v*_. In practice, we will use a relative threshold *t*_p_ ∈ (0, 1) to define *t*_*v*_ = *n t*_p_ – with *n* the number of amino acids.

We summarize the previous discussion with the following:

##### Definition. 1

*Using the filtration𝒢* _*u*_ *coding subsets of the path graph connecting the n amino acids of a polypeptide chain, we define:*

- *The* fraction *of positive critical points:* 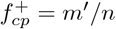.
- *The* mean persistence : 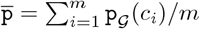,
- *The* normalized persistence entropy :

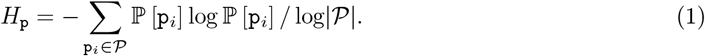
- *The maximum number of connected components observed:* 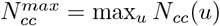

*Using the filtration H*_*v*_ *consisting of super-level sets of the function N*_*cc*_(*u*), *we define:*

- *The number of persistent local maxima of the function N*_*cc*_(*u*) *at persistence threshold t*_*v*_ : *PLM*(*t*_*v*_).

#### The pLDDT based filtration

The filtration𝒢 _pLDDT_ uses *−* pLDDT as parameter *u*. We indeed process *−*pLDDT values in increasing order, to handle high quality regions first. Assuming that all pLDDT values are different, the PD therefore contains exactly *n−* 1 critical points corresponding to the *n−* 1 edges connecting the *n* residues. A number of these have a null persistence and correspond to *accretion*–a given *C*_*α*_ carbon/amino acid inserted in the Union-Find data structure, and merged immediately to a neighbor to the left or the right. These account for the definition of the fraction 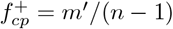 (Def. 1).

The c.c. appearing along the filtration 𝒢_pLDDT_ are units of the protein which form *independently* and eventually merge to compose the entire backbone. Phrased differently, the filtration provides a canonical way to find regions in the protein, making it possible to study their properties. In the sequel, we analyze these regions in terms of intrinsic disorder propensity as provided by `AIUPred` [17] and compliance with the null model presented in the next section.

### 2.2 Null model

In this section, we use the previous filtration to define a Doob martingale yielding a concentration result for the number of connected components.

We define a null model to assess the statistics of Def. 1. Consider a set of a.a. with consecutive indices. Intuitively, if these a.a. have coherent pLDDT values in some (narrow) interval, the rank of the a.a. as a function of their pLDDT value is expected to be a random permutation of the a.a. indices. We formalize this model and will use it as a yardstick while studying `AlphaFold2` predictions and the function 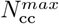.

#### Null model

Consider a fictitious protein with *n* a.a. having random pLDDT values uniformly distributed in fixed interval – that is [0, 100] for `AlphaFold2` predictions. This assumption is equivalent to saying that the ranks of the a.a. in the sorted sequence of pLDDT values is a random permutation *π* of [*n*]. After *k* insertions by increasing values, the occupied set is *S*_*k*_ = {*π*(1), …, *π*(*k*)}. We may also see the path graph as a string of 0 and 1s, with 1s corresponding to occupied positions. The number of connected components *C*_*k*_ after *k* insertions is then the number of runs of 1s.

From the combinatorial standpoint, we noted above that there are *n*(*n* + 1)*/*2 possible connected sub-graphs. Using probabilistic arguments, we further characterize our null model by the following theorem, whose proof relies on standard arguments, including the Azuma-Hoeffding inequality:

##### Theorem. 1

*The probability mass function of the number of c*.*c. C*_*k*_ *after k insertions is given by*

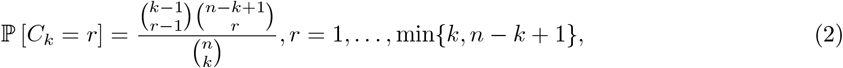

*and*

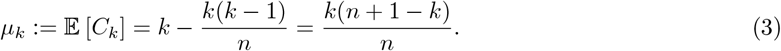

*The maximum of the expected number of connected components satisfies*

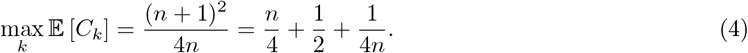

*Moreover*, 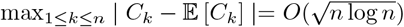 *with probability ≥* 1 *−* 1*/n*^2^.

The previous theorem is illustrated by plotting the curve of *C*_*k*_ as a function of *k*, or equivalently the function 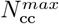as a function of the persistence parameter pLDDT (Fig. 1).

**Figure 1:**
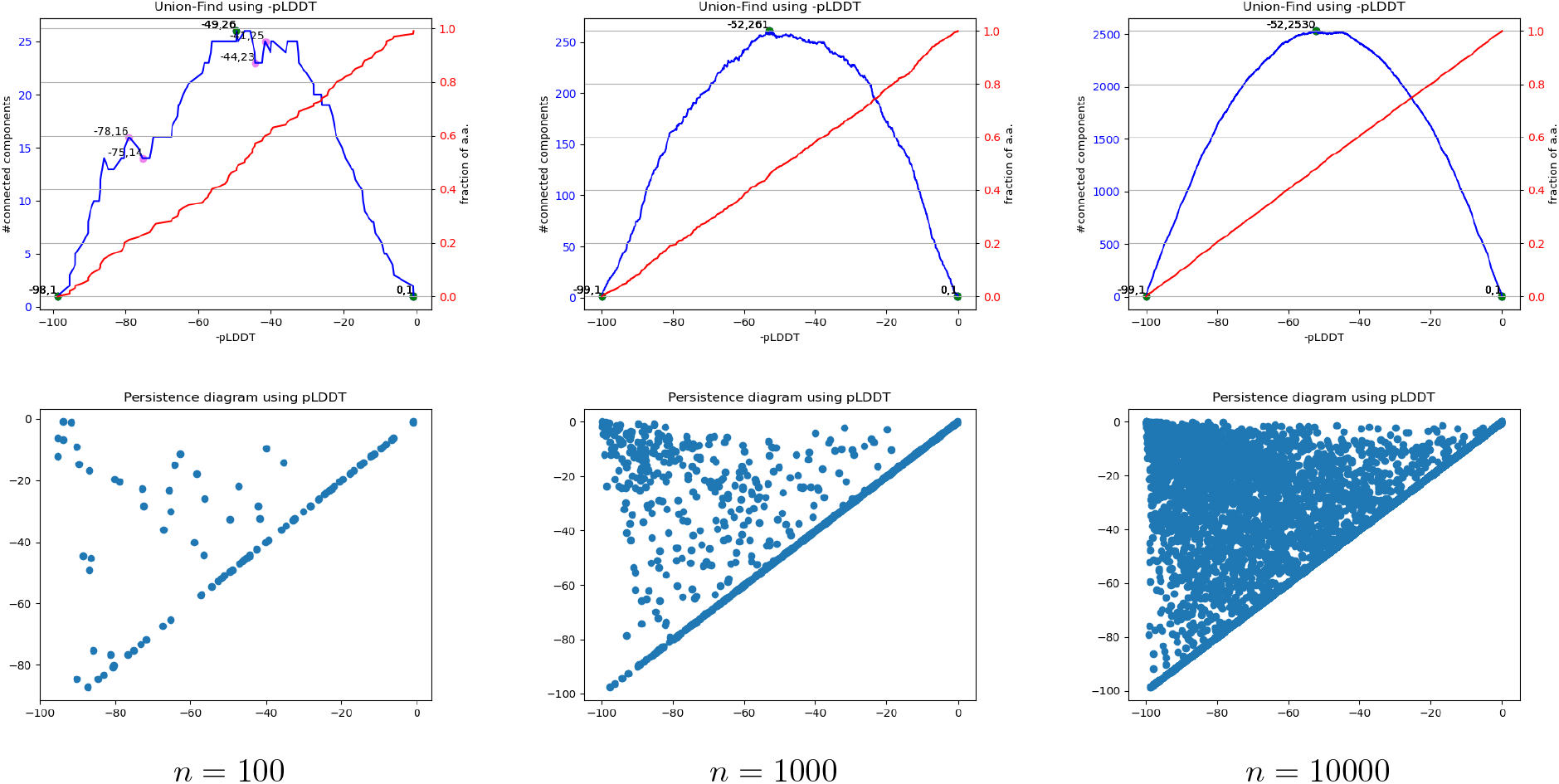
Filtration𝒢 _pLDDT_ on fictitious proteins with *n* nodes and uniformly random pLDDT values in [0, 100], obtained by inserting the amino acids by increasing value of*−* pLDDT. (Top row) Left scale and blue curve: evolution of the number of connected components *N*_cc_(*−* pLDDT). Right scale and red curve: cumulative fraction of amino acids (nodes). **(Bottom row)** Persistence diagrams for the number of connected components.

The persistence diagram also exhibits a characteristic triangular shape with a concentration of critical points near its apex, corresponding to the death of connected components upon moving forward to the full path graph in *𝒢*_pLDDT_. Simulations also give the following approximate values: 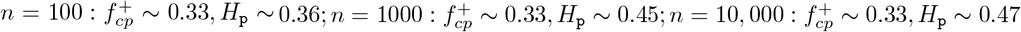.

##### Remark 1

*As an additional example, consider next the following fictitious protein–the so-called* stairs *model:(i) a first group of a*.*a. corresponding to an unstructured region on the N-ter side, with low confidence, say uniform pLDDT* ∈ [0, 49]; *and (ii) a second group of a*.*a. corresponding to a structured region on the C-ter side, with high confidence, say uniform pLDDT* ∈ [50, 100]. *Along the insertion, one observes the formation of the former and the latter regions (Fig. S2)*.

#### p-value calculations

We use theorem 1 to define p-values in several ways, so as to test *H*0 : *π∼* Uniform Permutation[n] versus *H*1 : *¬H*0.

If a value of *k* is fixed, say *k*_0_ = ⌊(*n* + 1)*/*2 and 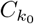 stands for the number of connected components for that value, we define:

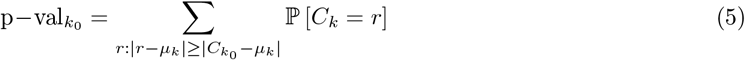

If one observes the whole sequence of insertions and the sequence of values, *C*_0_, …, *C*_*n*_ the derivation via Azuma’s bound (see theorem proof) gives a directly p-value. Since this is a conservative calculation (via upper-bounding) requiring large deviations to be significant, a more powerful test is obtained using a Monte Carlo / permutation test (PT) [18].

Let *T*_obs._ = max_*k*_ | *C*_*k*_*− µ*_*k*_ |the maximum value observed for the permutation of interest, and let *T*_*b*_ the corresponding test statistics for a random permutation *b* amidst *B* permutations: The following is the classical permutation p-value:

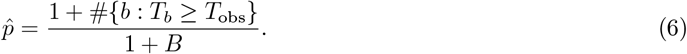

Note that the absolute value in the definition of *T*_obs._ counts deviations in both directions, that is, the test is two-sided.

Under H0, with *p*(*t*) = ℙ_*H*0_ [*T ≥ t*], the random variable *R* | *T*_obs._ = *t* follows a binomial distribution with parameters *B* and *p*(*t*), and Var [*R* | *T*_obs._ = *t*] = *Bp*(*t*)(1 *− p*(*t*)). Therefore

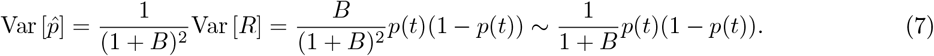

Imposing Var 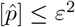 yields *B* ≥ *p*(1*− p*)*/ε*^2^ *−* 1. To control the Monte Carlo error at *p ∼ α* – *i*.*e*. near the decision boundary, we set *B* ≥*α*(1 *−α*)*/ε*^2^. For example, *α* = 0.05 and *ε* = *α/*10 yield *B∼* 1900. See Fig. S3.

#### Multiscale p-value calculations

The previous calculation takes into account the entire sequence of length *n*. But it can also be applied to any connected component formed during the insertion process. We noted above that there are *n*(*n* + 1)*/*2 such possible connected components, and for a particular insertion order, these are associated with the points of the persistence diagram. We therefore compute the pvalue for each such connected component, and report the scatter plot *connected component size* versus pvalue.

### 2.3 Software

The code to compute the statistics presented in this work is available within the Structural Bioinformatics Library [19]. See: [AlphaFold analysis](https://sbl.inria.fr/doc/Alphafold_analysis-user-manual.html), as well as [SBL portal](https://sbl.inria.fr/), [Documentation](https://sbl.inria.fr/doc/), [Applications](https://sbl.inria.fr/applications/), [Installation guide](https://sbl.inria.fr/doc/sbl-installation-guide.html).

## 3 Results

We use the filtration𝒢 _pLDDT_ and the associated values (Def. 1, Fig. 2, Fig. 4, Fig. 5) to characterize the coherence of pLDDT values along the backbone and the fragmentation of `AlphaFold2` reconstructions.

**Figure 2:**
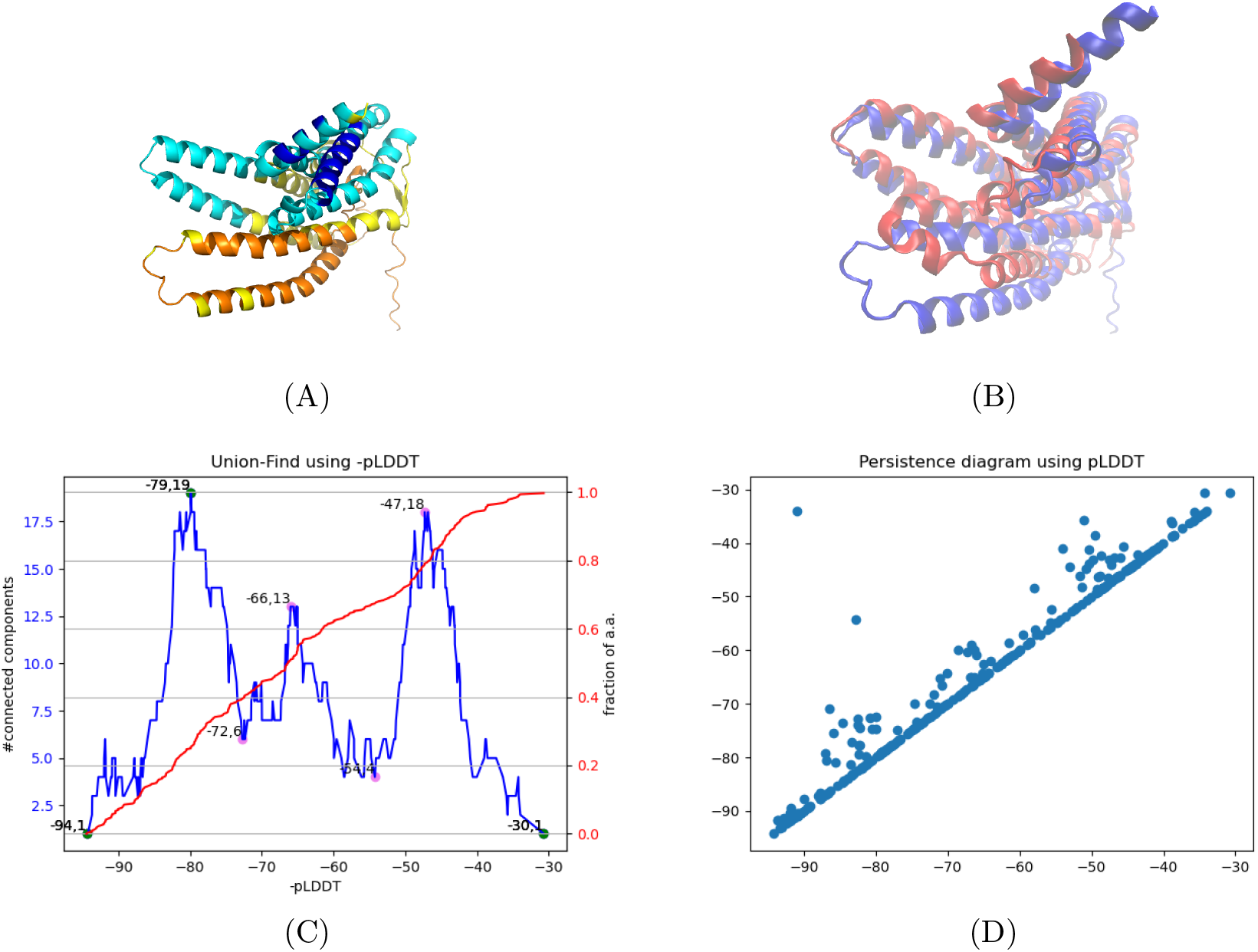
Human olfactory receptor: model AF-Q9H342-F1-model_v4 versus chain R of the experimental structure 8hti: persistent maxima of the curve *N*_cc_(*−* pLDDT) counting connected components and regions with loose structural alignment against the PDB structure. **(A)** `AF-Q9H342-F1-model_v4.pdb` models the human olfactory receptor 51J1 – 316 amino acids. **(B)** Structural alignment with the homolog chain R from `8hti.pdb` (E-Value: 1.943e-62), yielding a *lRMSD* = 6.12 for 303 aligned residues. **(C)** The function *N*_cc_(*−*pLDDT) yields three persistent local maxima characterized by *N*_cc_(*−*79) = 19, *N*_cc_(*−*66) = 13 and *N*_cc_(*−*47) = 18. **(D)** Persistence diagram.

**Figure 3:**
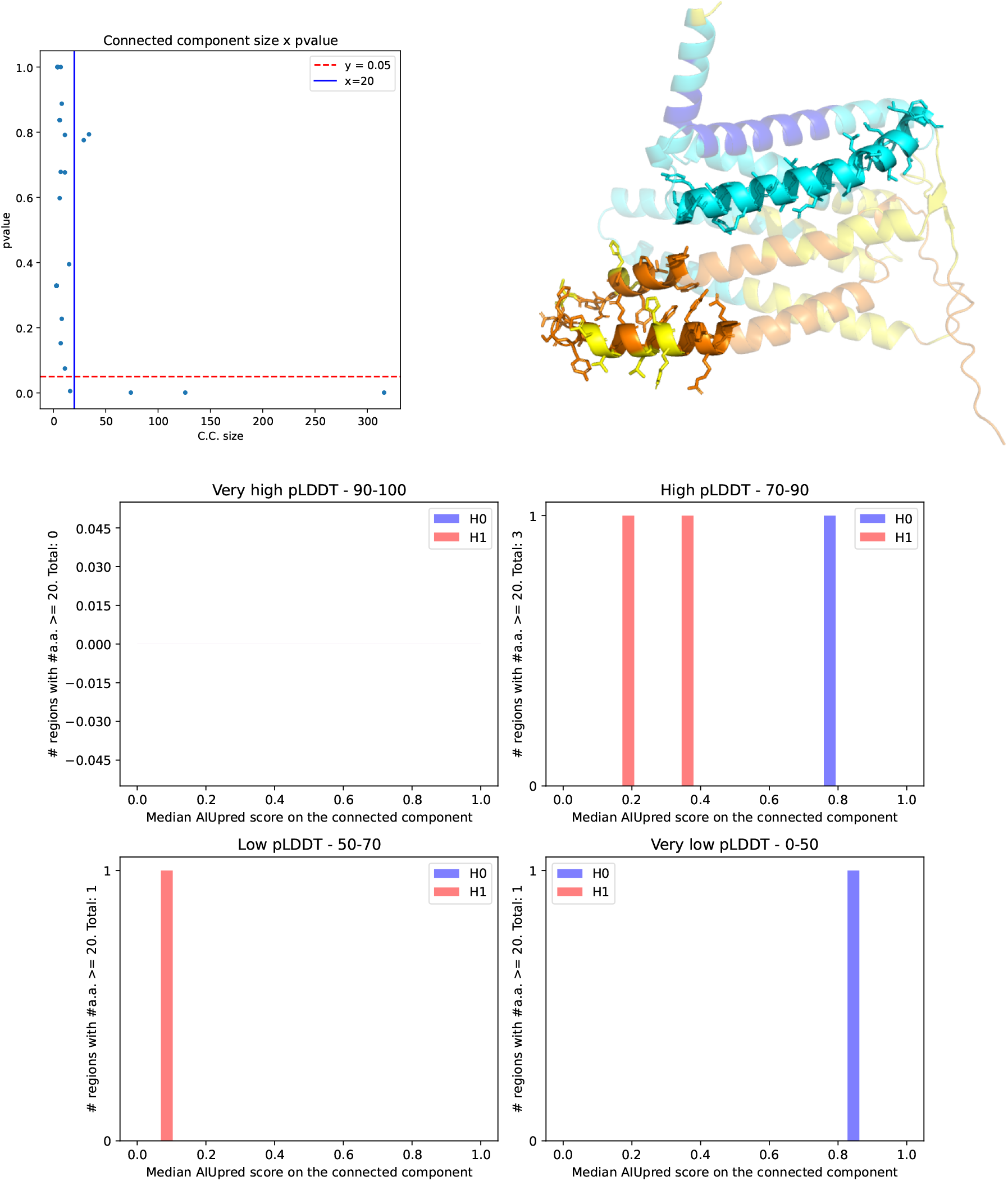
Human olfactory receptor. Polyptych plot with minimum connected component size of 20. (First row) Scatter plot for pvalues and structure. Regions with sticks correspond to c.c. marked with a * in Table 1. Note the correspond to the a.a. ranges 62-90 and 121-154 respectively. **(Second row)** Polyptych plot.

**Figure 4:**
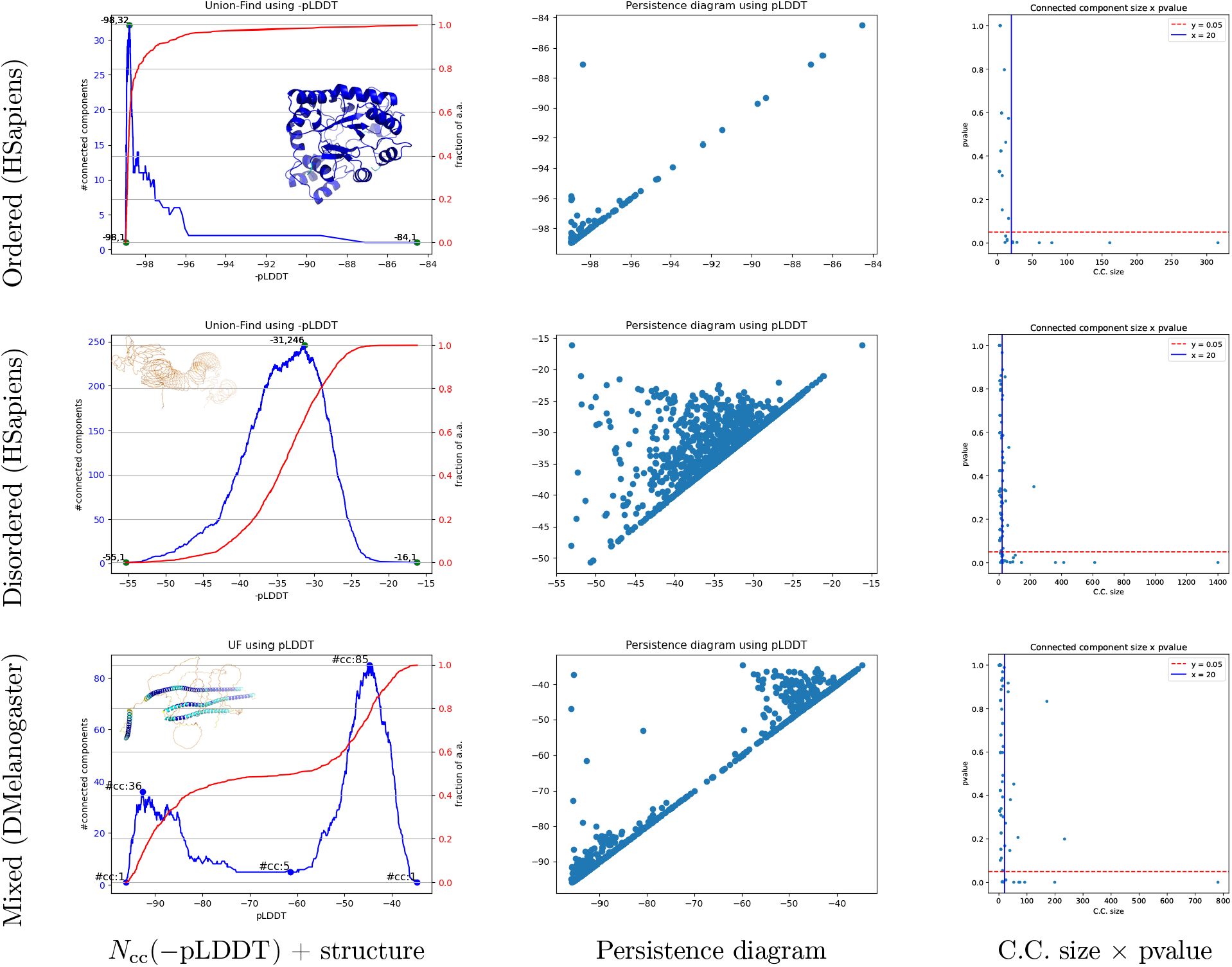
Filtrations𝒢 _pLDDT_ on three prototypical examples corresponding to ordered, disordered and mixed proteins: evolution of the number of connected components *N*_cc_(*−*pLDDT), persistence diagram, and pvalues. **(First row)** (H. Sapiens; AF-P15121-F1-model_v4; 316 amino acids; *H*_p_ *∼* 0.04.) **(Second row)** (H. Sapiens; AF-Q02817-F15-model_v4; 1400 amino acids; *H*_p_ *∼* 0.36.) **(Third row)** (D. Melanogaster; AF-Q9VQS4-F1-model_v4; 781 amino acids; *H*_p_ *∼* 0.28.)

**Figure 5:**
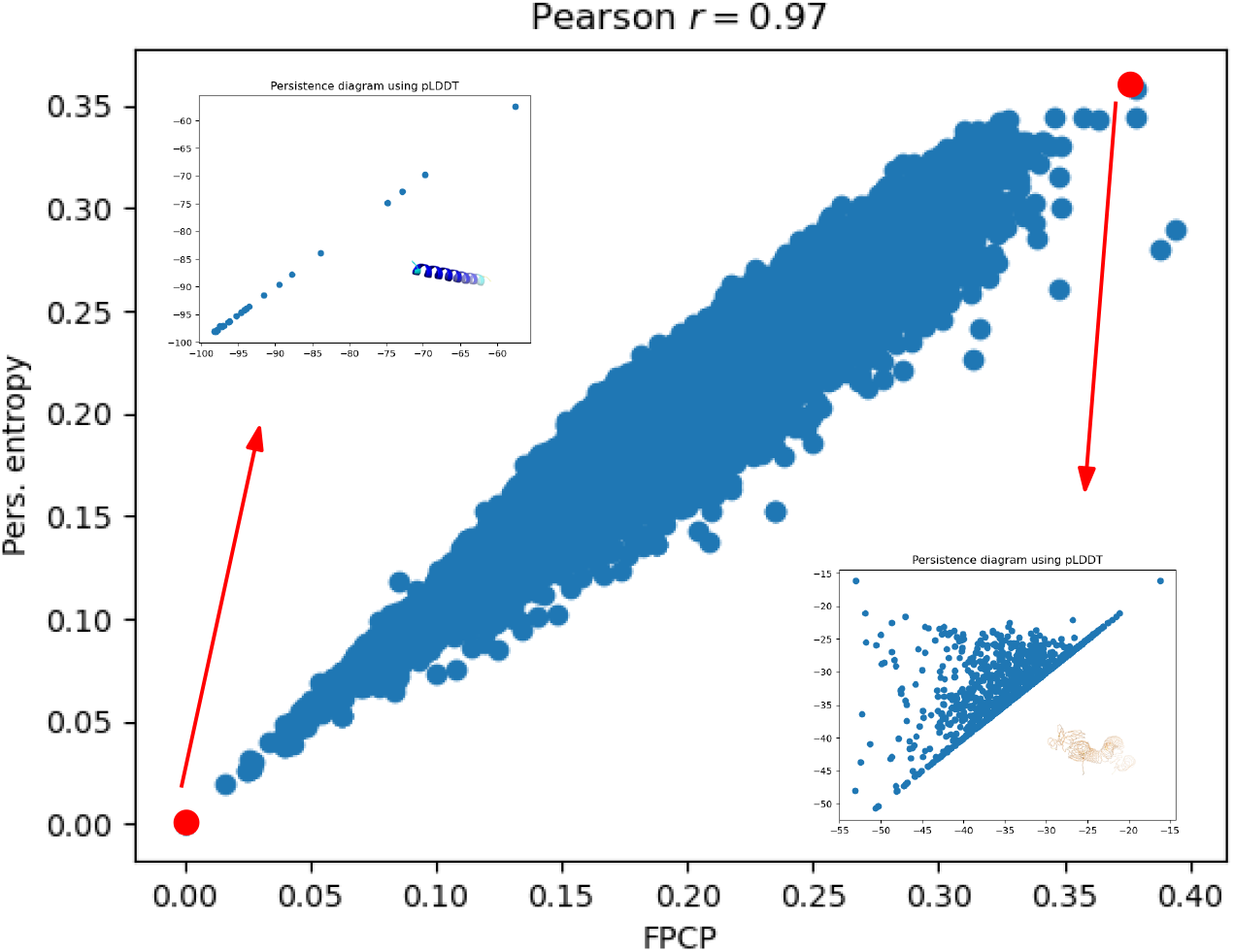
Homo Sapiens genome wide scale: persistence entropy *H*_p_ vs fraction of positive critical points (FPCP) 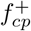 of 𝒢_pLDDT_ show a continuum between highly coherent and random-like structures in terms of pLDDT. Insets displaying the structures achieving the minimum and maximum *H*_p_ values. The corresponding models are AF-O00631-F1-model_v4 (*H*_p_ = 0) and AF-Q02817-F15-model_v4 (*H*_p_ = 0.36).

We start with one example, and proceed with a systematic analysis of statistics defined in Def. 1.

### Fragmentation, a case study: the human olfactory receptor 51J1

We consider the human olfactory receptor 51J1 (316 amino acids), file `AF-Q9H342-F1-model_v4.pdb`. A sequence based search in the PDB provides as homolog chain R from the experimental structure `8hti.pdb`, with the following statistics: Sequence Identity: 38%, E-Value: 1.943e-62, Region: 13-323. A structural alignment between the `AlphaFold2` model and chain R yields a *lRMSD* = 6.12, based on 303 aligned residues (Fig. 2). The curve *N*_cc_(*−* pLDDT) clearly identifies three persistent local maxima corresponding to pLDDT values of 47 and 79 respectively (Fig. 2 for the first and the third). The shape of function *N*_cc_(*−* pLDDT) around these three maxima is reminiscent from that of the null model. Yet, the null model is statistically stringent, and the calculation of pvalues for all connected components identified from the persistence diagram show that only two c.c. of size *>* 20 are consistent with this null model – that is one does not reject H0. As expected,these two c.c. are subsets from the regions (attraction basins) associated with the minima at pLDDT = 47 and pLDDT = 79 respectively. Interestingly, the region tagged (2H) is that where the AF model departs significantly from the experimental structure (Fig. 2).

As opposed to the classical AlphaFold labels *very high, high, low, very low*, our analysis identifies coherent regions in terms of pLDDT values and further characterizes their compliance with the null model.

### Different profiles revealed

Further insights are provided by the analysis of the function *N*_cc_(*−* pLDDT) and the persistence entropy *H*_p_, for various protein profiles, namely well ordered, disordered, and mixed (Fig. 4). It appears that the function *N*_cc_(*−*pLDDT) varies from unimodal (when considering persistent maxima) to multimodal. The former occurs for well structured (ordered) or unstructured proteins and the latter for mixed cases (Fig. 4). For example, the *D. Melanogaster* protein AF-Q9VQS4-F1-model_v4 exhibits a clear bimodal profile with two persistent maxima (Fig. 4). On this example, the plateau at five connected components corresponds to five helices–four long ones plus a shorter one. The persistence diagram associated with the construction of the filtration also exhibit the dynamics of connected components (Fig. 4). The PD of the human protein AF-Q02817-F15-model_v4 has the typical triangular profile observed for the null random model. The PD of *D. Melanogaster* quantifies the dynamics of the high (resp. low) confidence region of the molecule on the left (resp. right) part of the diagram. We note that the apex of the triangles are not as populated as those of the null model (Fig. 1), showing a lesser randomness of pLDDT values, which is consistent with the lesser entropy *H*_p_.

The scatter plots of pvalues computed for all c.c. associated with points of the persistence diagram (Fig. 4) enable identifying connected regions complying with the null model. While many small components (say size *<* 20 amino acids) complying with the null model are observed in all cases, only proteins with *disordered* regions exhibit large components, *>* 200 a.a. for AF-Q02817-F15-model_v4 and AF-Q9VQS4-F1-model_v4 (See also SI Tables of p-values for connected components S2, S3, S4.)

### Persistence entropy at the genome wide scale

At the human genome wide scale, we observe a strong correlation between the three pairs of variables amidst 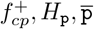 (Fig. 5, Fig. S4). The maxima of 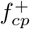 are consistent with those a the null random model, while those of *H*_p_ are slightly below those of the same null model. (NB: average values of 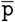 are irrelevant for the null model.) For all organisms, the minimum *H*_p_ is observed for very regular structures whose pLDDT values are within a small range (Fig. S5. Instead, the maxima are observed for highly disordered structures whose a.a. span a wide range of pLDDT values (Fig. S5). For such cases, the typical triangular shape in the PD is also observed.

These statistics indicate that all predictions form a continuous spectrum ranging from highly coherent to random-like structures.

### Maximum value 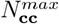of the function *N*_cc_(*−*pLDDT)

The maximum number of local maxima observed 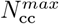 normalized by the protein size (Fig. S6) in general lower than the expected maximum 1*/*4, even though some cases reach or even surpass that bound. This is accounted for by two reasons. First, accretion is at play–see definition above. Second, pLDDT values come in batches, so that local insertions into *stretches* along the sequence prevent the maximum number of local maxima to rise. These batches are identified by the persistent local maxima discussed next.

### Persistent local maxima PLM of the function *N*_cc_(*−* pLDDT)

We first note that the number of persistent local maxima is in general less than 10 (Fig. S4). Also, the correlation between *H*_p_ and PLM is quite low (Fig. S4), which is expected. One can indeed have a significant number of persistent critical points localized in a narrow range of pLDDT values (Fig. S7). This phenomenon is well known to geographers defining peaks on mountains: two tall peaks can be separated by a low lying pass, but be located at a short flying distance [20]. Nevertheless, as pointed out above, persistent local maxima identify coherent stretches along the sequence in terms of pLDDT values.

### Model assessment and database queries

In the model assessment perspective, querying the encoding provided by the number of a.a., the persistence entropy *H*_p_, the fraction of positive critical points 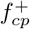 and the number of persistent local maxima PLM enables the identification of cases incurring a large fragmentation. Such cases exhibit a wide range of pLDDT values aggregated in coherent stretches along the sequence, yielding a large entropy and a (relatively) high number of persistent local maxima of the function *N*_cc_(*−* pLDDT) (Fig. 6).

**Figure 6:**
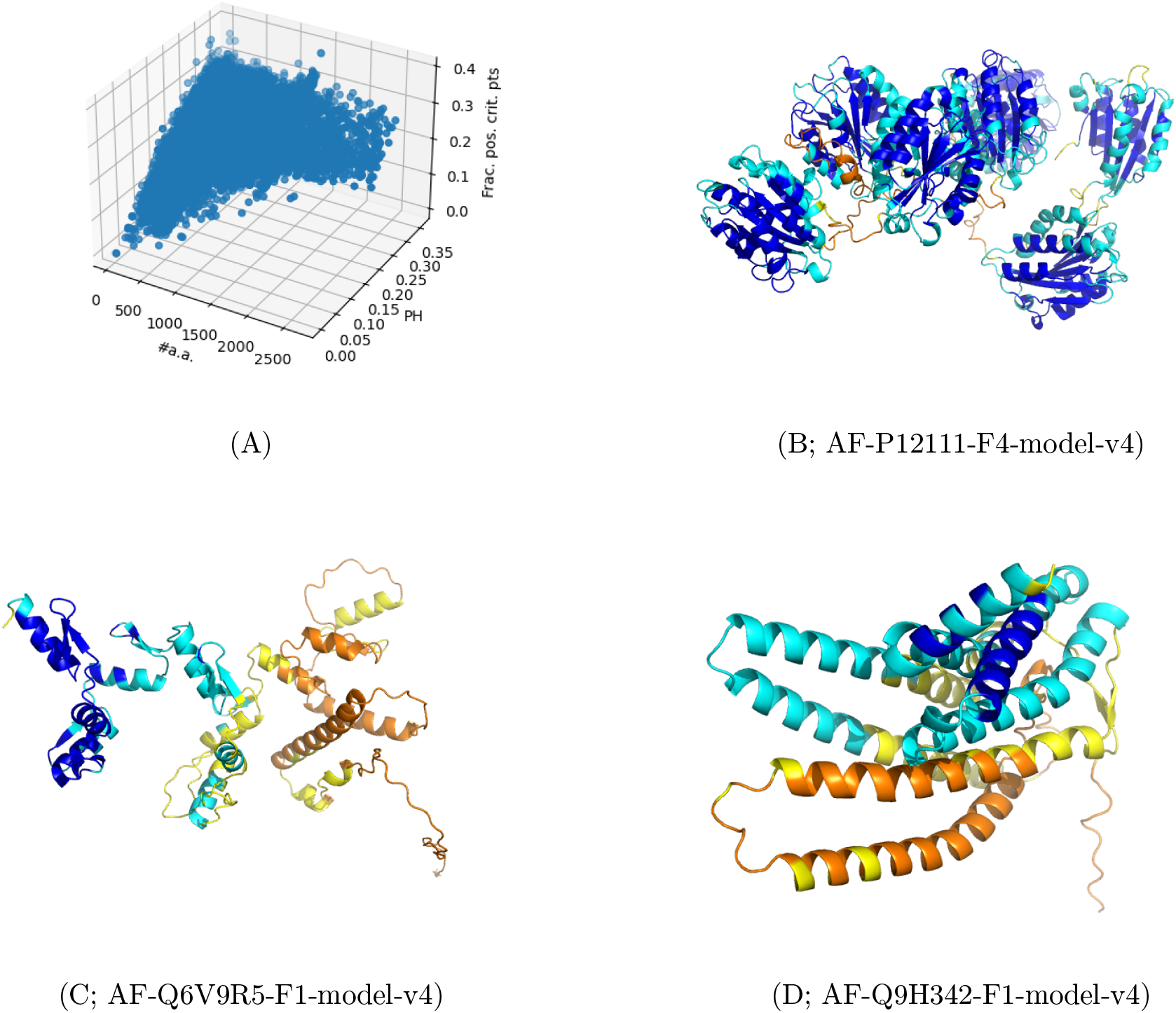
Homo Sapiens predictions. Illustration of model fragmentation: wide range of pLDDT values aggregated in coherent stretches of pLDDT values along the sequence. **(A)** Scatter plot with protein size*×* persistence entropy *H*_p_ *×*fraction of positive critical points 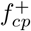. **(B, C, D)** At persistence threshold *t*_p_ = 0.025, 86 structures are characterized by *H*_p_ ≥0.25, #*a*.*a*. ≥200, PLM≥ 3. Three of them are displayed.

The ability to identify such cases is of high interest to further our understanding of the learning process implemented by recent deep learning based structure predictors.

### Connected components: pLDDT values, intrinsic disorder, and null model

As noticed earlier, the c.c. seen along the filtration can be used to study specific properties. Of particular interest is the disordered nature of individual residues, which we evaluate using the `AIUPred` software [17]. We summarize this prediction on a region by the median `AIUPred` value of its amino acids.

For a protein, consider all c.c. larger than a user specified threshold, *e*.*g*. 20 residues. We ascribe these c.c. to four buckets corresponding to the four usual pLDDT ranges, namely very low, low, high, very high.

We then plot the histogram of `AIUPred` median values (one per c.c.), which we further paint in blue when the c.c. complies with H0 of our null model, and red otherwise. We term the resulting plot the *polyptych plot* (Fig. 3). This plot has two interests:

- (i) For a quadrant, irrespective of the H0/H1 tag, discrepancies between the pLDDT values and the `AIUPred` scores are evident. In particular, regions with low (resp. high) pLDDT values and low (resp. high) `AIUPred` scores call for an inspection.
- (ii) Within a quadrant, the H0 and H1 tags are of further interest. In particular, accepting H0 for a relatively narrow range of pLDDT scores is a very strong evidence of the homogeneity of `AlphaFold2`’s (lack of) confidence for that region.

As an illustration, consider the olfactory receptor (Fig. 2, Fig. 3, Table 1). Using a minimum c.c. size of 20, it appears that two regions out of seven qualify with H0, with statistical summary (#a.a.: 29; pLDDT score: 78.62; `AIUPred` score: 0.77), and (#a.a.: 34; pLDDT score: 46.93; `AIUPred` score: 0.85) (Fig. 3). With a high pLDDT median value and a high AIUPred median score, the first one lacks coherence. With low pLDDT and AIUPred statistics, the second one appears coherently suspicious (Fig. 3).

As a final case, one may consider the region associated with low pLDDT, whose signature is (#a.a.: 316; pLDDT score: 66.41; `AIUPred` score: 0.09). The low pLDDT and AIUPred values are not problematic in this case, since this region actually corresponds to the entire sequence–spanning the large pLDDT range [30, 94].

Additional examples are provided in SI (Fig. S2, Fig. S3,Fig. S4). In these, we have have systematically highlighted regions characterized by (i) high or very high median pLDDT score, and (ii) median `AIUPred` score *≥*.5.

## 4 Outlook

Quality assessment is a key step in leveraging the results of deep learning based structure prediction methods in general, and those of `AlphaFold2` in particular. This work introduces novel tools to identify stretches of amino acids displaying *coherent* pLDDT values.

The first level of our analysis consists of identifying stretches of residues belonging to the *attraction basins* of persistent local maxima. The second one hinges on a combinatorial notion of coherence stating that within a stretch, the pLDDT scores are statistically indistinguishable from one another, as if drawn from a uniformly random distribution.

Using the human genome of the AlphaFold Protein Structure Database, we show that predictions face a whole array of situations, from entirely coherent proteins, to proteins split into several distinct coherent regions.

The resulting partitioning is directly relevant for the study of various biophysical properties, including the annotation of functional regions and the relationship to disorder. On a more fundamental agenda, the regions identified offer a new perspective to understand the learning biases inherent to deep learning based structure prediction tools.

## 5 Artwork

**Table of p-values for connected components 1** Input file: AF-Q9H342-F1-model_v4

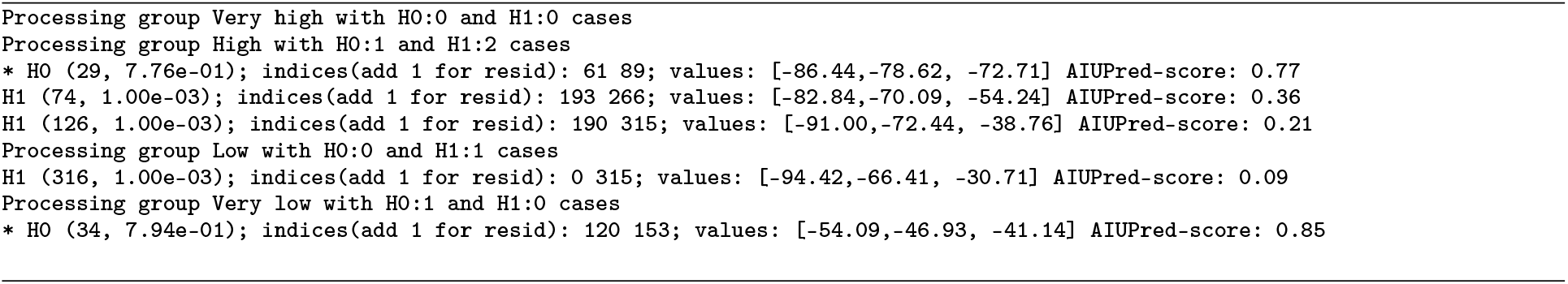

## Acknowledgments

This work has been supported by the French government, through the 3IA Côte d’Azur Investments (ANR-19-P3IA-0002), and the ANR project Innuendo (ANR-23-CE45-0019).

Nicolas Broutin, Vincent Mallet, and the reviewers are acknowledged for stimulating discussions.

## Data availability

This manuscript is not associated to any new data.

The code to compute the statistics presented in this work is available within the the Structural Bioinformatics Library [19]. See: <u>AlphaFold analysis</u>, as well as <u>SBL portal</u>, <u>Documentation</u>, <u>Applications</u>, <u>Installation guide</u>.

