## Supplementary material for "Characterizing the fragmentation of AlphaFold predictions": SI

# S1 SI

### S1.1 pLDDT statistics on whole genomes

| genome / pLDDT | 50 | 70 | 90 |
|---|---|---|---|
| AThaliana/all | 0.204 | 0.315 | 0.550 |
| CAlbicans/all | 0.213 | 0.312 | 0.570 |
| CElegans/all | 0.202 | 0.323 | 0.597 |
| DDiscoideum/all | **0.288** | 0.423 | **0.681** |
| DMelanogaster/all | 0.281 | 0.387 | 0.623 |
| DRerio/all | 0.246 | 0.343 | 0.593 |
| EColi/all | **0.029** | **0.078** | 0.275 |
| GMax/all | 0.217 | 0.338 | 0.577 |
| HSapiens/all | 0.284 | 0.382 | 0.666 |
| MJannaschii/all | 0.035 | 0.084 | **0.264** |
| MMusculus/all | 0.255 | 0.352 | 0.597 |
| OryzaSativa/all | 0.257 | **0.408** | 0.623 |
| RattusNorvegicus/all | 0.253 | 0.351 | 0.596 |
| SCerevisiae/all | 0.213 | 0.314 | 0.582 |
| SPombe/all | 0.187 | 0.289 | 0.558 |
| ZeaMays/all | 0.260 | 0.394 | 0.635 |
| SAureus/all | 0.045 | 0.096 | 0.295 |
| HPylori/all | 0.053 | 0.123 | 0.347 |
| MTuberculosis/all | 0.069 | 0.133 | 0.322 |
| Aeruginosa/all | **0.036** | **0.086** | **0.278** |
| PFalciparum/all | **0.460** | **0.584** | **0.804** |

Table S1: **Cumulated distribution function for pLDDT values of all a.a. of all proteins in a genome.** Top: model genomes. Bottom: five *global health* proteomes. Min and max values per column stressed in bold.

### S1.2 Algorithms

### S1.3 Proof of theorem 1

*Proof.*

**Exact distribution – Equation 2.** After $k$ insertions, the number $r$ of c.c. lies in the range $1, \ldots, \min\{k, n-k+1\}$. For the upper bound, note that there are $n-k$ free slots which determine at most $n-k+1$ runs, so that the minimum between this value and $k$ is taken.

To derive the exact probability of $C_k$ assuming $r$ runs, note the following:

- The lengths $L_i$ of the $r$ runs satisfy $L_1 + \cdots + L_r = k$ with $L_i \geq 1$. The number of solutions is given by the number of compositions of $k$ into $r$ parts, namely $\binom{k-1}{r-1}$.
- Placing $r$ runs into $n-k+1$ slots yields $\binom{n-k+1}{r}$ choices.

Putting together yields Eq. 2.

**procedure** BUILD_PATH_GRAPH_FILTRATION($[(j, u_j)]_{i=1,\ldots,n}$)
    Form the list $[(j, u_j)], j = 1, \ldots, n$, for the $n$ amino acids
    Let $L$ be this sorted list ascending $u_j$ values
    **for** $(j, u_j) \in L$ **do**
        UF.make_set($c_j$)
        if $j > 1$ and $c_{j-1}$ exists in the UF data structure: UF.union($c_j, c_{j-1}$)
        if $j < n$ and $c_{j+1}$ exists in the UF data structure: UF.union($c_j, c_{j+1}$)
        $cc_j \leftarrow$ UF.num_cc
        $nn_j \leftarrow nn_j + 1$
    **end for**
    **return** $\{(cc_i, nn_i)\}$ and the associated persistence diagram
**end procedure**

Figure S1: **Union-Find on the polypeptide chain and filtration $\mathcal{G}_u$.** Particular case: using pLDDT as value for the parameter $u$ yields the filtration $\mathcal{G}_{\text{pLDDT}}$.

**Maximum – Equation 4.** After $k$ insertions, one has $C_k = k - A_k$, with $A_k$ the number of edges $(i, i+1)$ of the path graph whose two endpoints are occupied by a 1. In the case of random insertions, the probability that both endpoints of a given edge are occupied is equal to

$$\frac{k}{n} \frac{k-1}{n-1},$$

and we get

$$\mathbb{E}[A_k] = (n-1)\frac{k}{n}\frac{k-1}{n-1} = \frac{k(k-1)}{n}.$$

This yields

$$\mathbb{E}[C_k] = k - \frac{k(k-1)}{n} = \frac{k(n+1-k)}{n}. \tag{8}$$

The maximum of this function is attained at $k^* = (n+1)/2$, whence Eq. 4.

**Concentration bound.** We wish to know how $\max_k C_k$ is concentrated near its expectation.

Since we insert nodes one by one, consider the following filtration of $\sigma$-algebras:

$$\mathcal{F}_i = \sigma(\pi(1), \ldots, \pi(i)), i = 0, \ldots, k. \tag{9}$$

We use this filtration to define the following Doob Martingale [1] for $C_k$:

$$M_i := \mathbb{E}[C_k | \mathcal{F}_i], i = 0, \ldots, k. \tag{10}$$

We have $M_0 = \mathbb{E}[C_k]$ – no conditioning, and $M_k = C_k$ – complete information available.

Upon adding the node $\pi(i)$, $\Delta C_k \in \{-1, 0, 1\}$, which corresponds to the merge of two c.c., the accretion of a node to an existing c.c., or the creation of a new c.c. The insertion of $\pi(i)$ admits $\binom{n-(i-1)}{k-(i-1)}$ equally likely possibilities. The calculation of $M_{i-1}$ involves all scenarios – those in which $\pi(i)$ is revealed and those in which it is not. The calculation of $M_i$ involves the scenarios in which $\pi(i)$ is involved only, with $\mid \Delta C_k \mid \leq 1$. Therefore the martingale increments satisfy deterministically

$$\mid M_i - M_{i-1} \mid \leq 1, \forall i \leq k. \tag{11}$$

[1] Note that the martingale property is satisfied due to the tower property of conditional expectations: $\mathbb{E}[M_{i+1}|\mathcal{F}_i] = \mathbb{E}[\mathbb{E}[C_k|\mathcal{F}_{i+1}]|\mathcal{F}_i] = \mathbb{E}[C_k|\mathcal{F}_i] = M_i$.

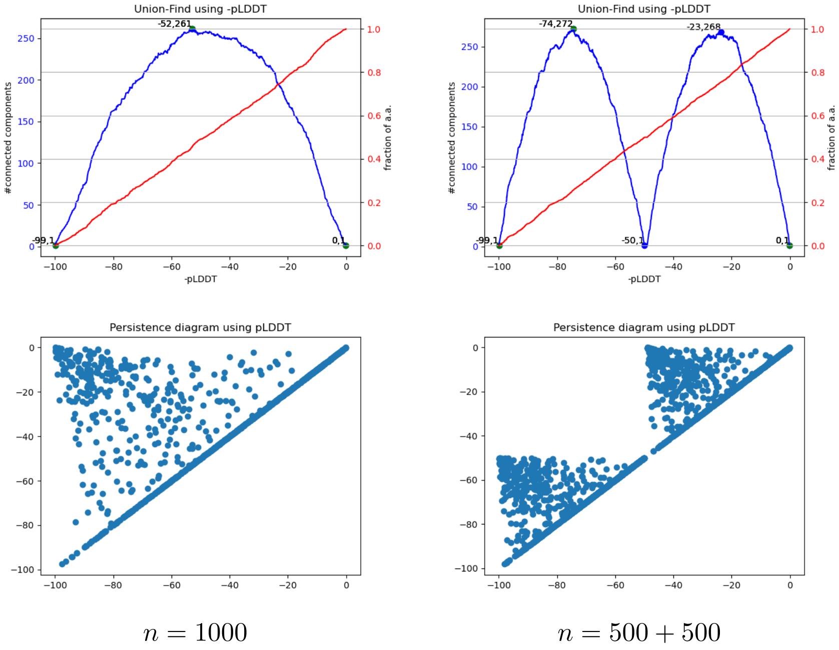

Figure S2: **Filtration $\mathcal{G}_{\mathbf{pLDDT}}$: comparison of the *flat* and *stairs* models.** Left: the $n = 1000$ a.a. are assigned random pLDDT values in $[0, 100]$; Right: the first 500 a.a. are assigned random pLDDT values in $[0, 49]$, and the last 500 a.a. are assigned random pLDDT values in $[50, 100]$. **(Top row)** Blue curve: function $N_{\text{cc}}(\text{pLDDT})$; red curve: fraction of amino acids. **(Bottom row)** persistence diagram.

The Azuma-Hoeffding inequality [21, 22] yields the following concentration inequality at each $k$:

$$\mathbb{P}\left[\mid C_k - \mathbb{E}\left[C_k\right] \mid \geq t\right] = \mathbb{P}\left[\mid M_k - M_0 \mid \geq t\right] \leq 2\exp\left(-\frac{t^2}{2\sum_{i=1,\ldots,k} 1^2}\right) = 2\exp\left(-\frac{t^2}{2k}\right). \tag{12}$$

To control the bound on the entire insertion process, we union-bound the previous expression:

$$\mathbb{P}\left[\max_{1\leq k\leq n} \mid C_k - \mathbb{E}\left[C_k\right] \mid \geq t\right] \leq \sum_k \mathbb{P}\left[\mid C_k - \mathbb{E}\left[C_k\right] \mid \geq t\right] \leq \sum_{k=1}^{n} 2\exp\left(-\frac{t^2}{2k}\right) \leq 2n\exp(-t^2/(2n)). \tag{13}$$

We get

$$2n\exp(-t^2/(2n)) \leq \delta \Rightarrow t \geq \sqrt{2n(\log(2n) - \log\delta)}. \tag{14}$$

Taking $\delta = 1/n^2$ yields $t^2 = 2n(\log 2 + 3\log n)$ and $\max_{1\leq k\leq n} \mid C_k - \mathbb{E}\left[C_k\right] \mid = O(\sqrt{n\log n})$ with probability $\geq 1 - 1/n^2$. □

### S1.4 Results–general

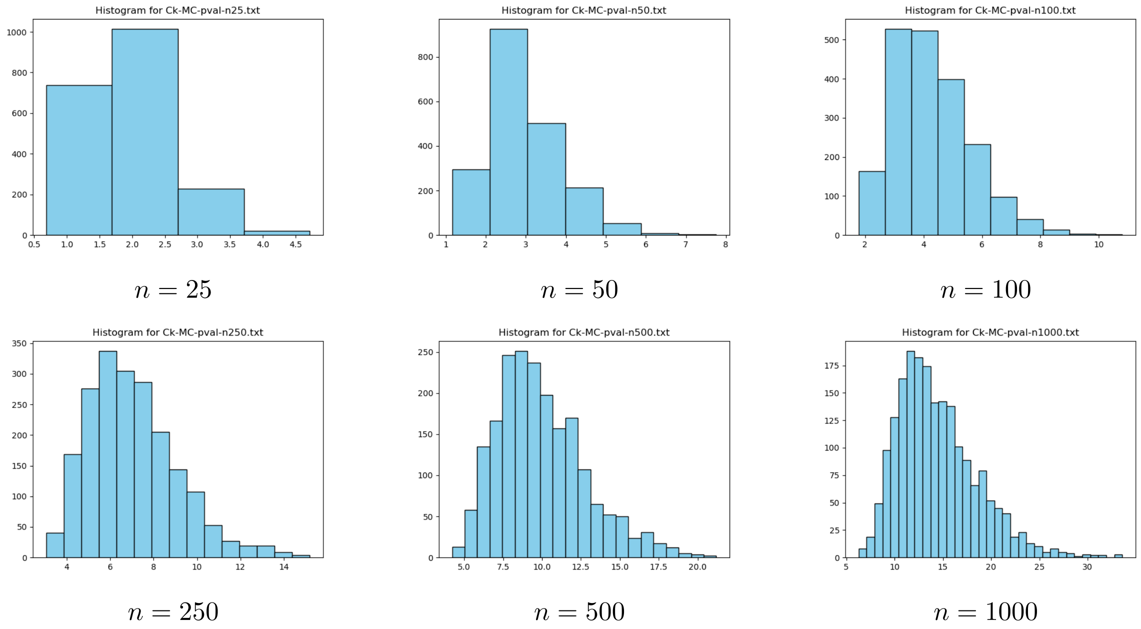

Figure S3: **Distribution of the test statistic $T = \max_k \mid C_k - \mu_k \mid$ for various values of $n$.** $B = 2000$ random permutations used in each case. Distributions used to compute the permutation p-value of Eq. 6.

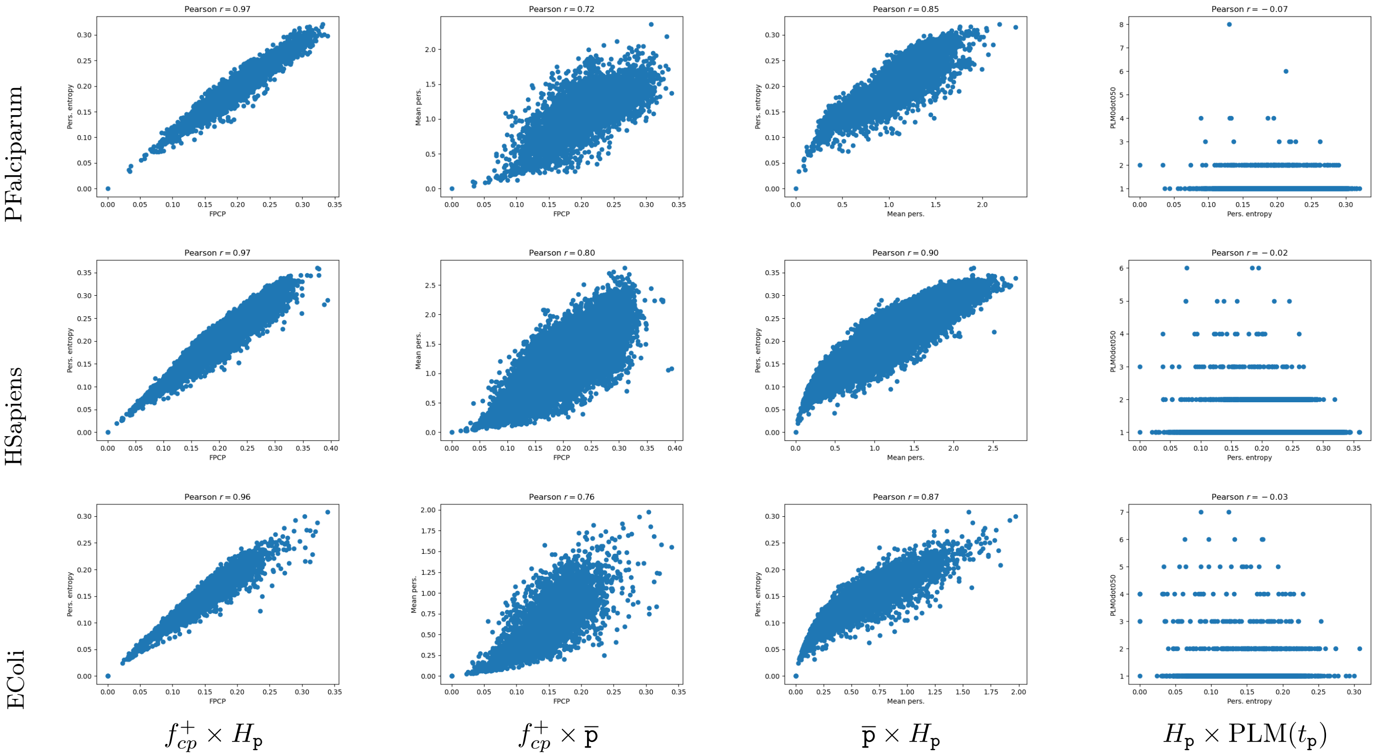

Figure S4: **PAeruginosa, HSapiens, Ecoli: correlations between the fraction of positive critical points $f_{cp}^{+}$, the persistence entropy $H_{\mathrm{p}}$, the mean persistence $\overline{\mathrm{p}}$, and the number of persistent local maxima PLM($t_{\mathrm{p}} = 0.05$).** Along these three examples: the fraction of positive critical points increases and so does the persistence entropy. Compare with Table S1.

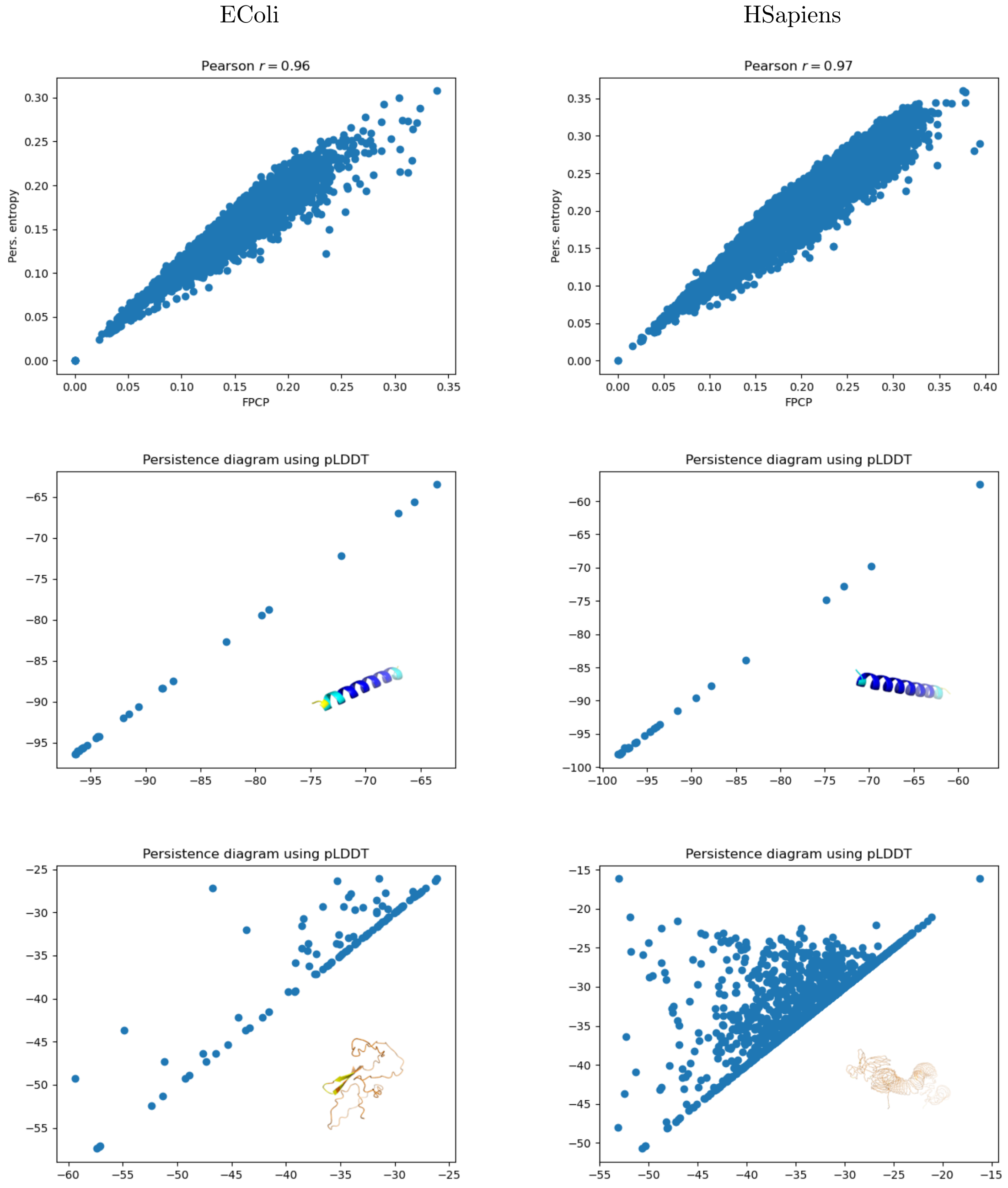

Figure S5: **Persistence entropy $H_{\text{p}}$ and fraction of positive critical points (FPCP) $f^{+}_{cp}$: illustrations on two genomes, with structures achieving the minimum and maximum values.** Compare with Table S1. **(First row)** Scatter plot fraction of positive critical points $f^{+}_{cp}$ $\times$ persistence entropy $H_{\text{p}}$. **(Second row)** Per organism, persistence diagram and `AlphaFold2` model achieving the minimum entropy $H_{\text{p}}$. **(Third row)** Per organism, persistence diagram and `AlphaFold2` model achieving the maximum entropy $H_{\text{p}}$. **(Structures for EColi)** AF-A5A617-F1-model_v4, 27 a.a.; AF-P62066-F1-model_v4, 106 a.a. **(Structures for HSapiens)** AF-O00631-F1-model_v4, 31 a.a.; AF-Q02817-F15-model_v4, 1400 a.a.

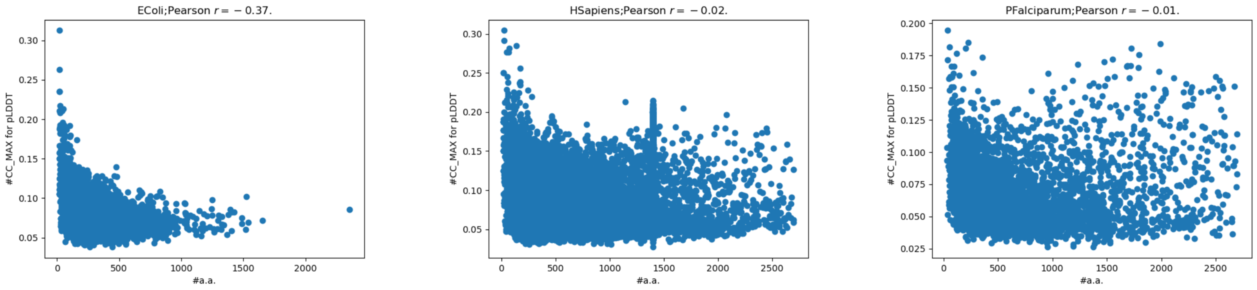

EColi HSapiens PFalciparum

Figure S6: **Protein size $n$ vs ratio $N_{\mathbf{cc}}^{max}/n$: illustration for three genomes.** By Theorem 1, the maximum expected value for $N_{\text{cc}}^{max}/n$ is 1/4.

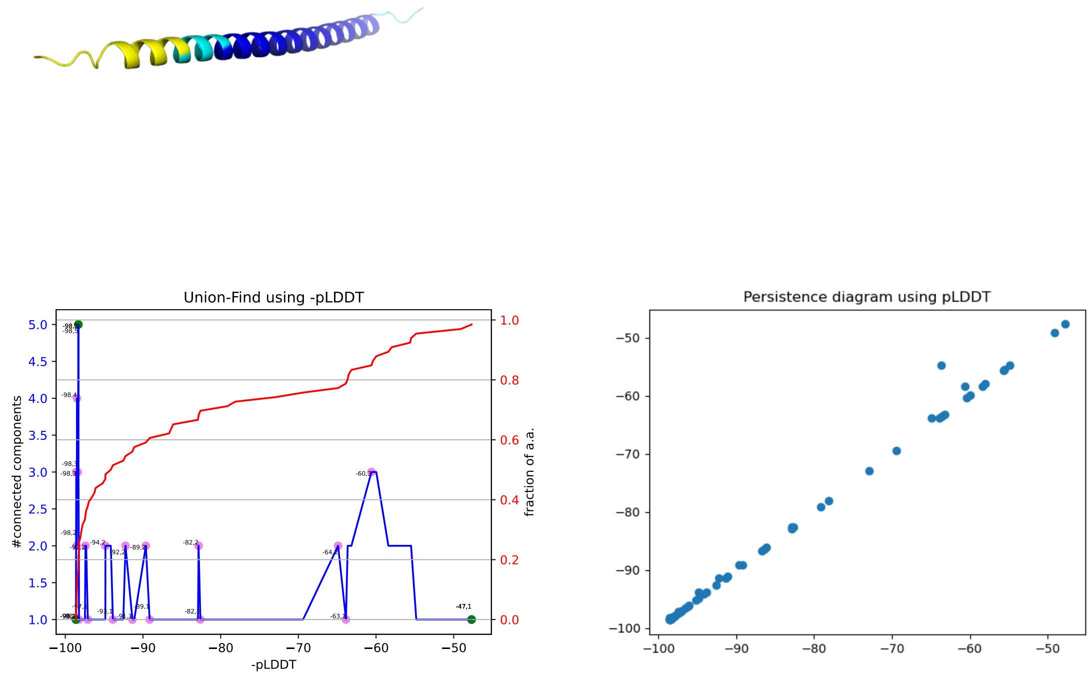

Figure S7: **Function $N_{\mathbf{cc}}(\mathbf{pLDDT})$ and its maximum value $N_{\mathbf{cc}}^{max}$ versus persistent local maxima PLM.** Illustration with AF-Q8N300-F1-model_v4
.

**Table of p-values for connected components S2** Input file: AF-P15121-F1-model_v4.pdb

```
sbl-alphafold-dbrun.py --pval-cc-size 20 --aiu  -f human-AF-P15121-F1-model_v4.pdb
...
Processing group Very high with H0:0 and H1:7 cases
H1 (28, 3.00e-03); indices(add 1 for resid): 175 202; values: [-98.95,-98.85, -98.74]AIUPred-score: 0.46
* H1 (22, 1.00e-03); indices(add 1 for resid): 48 69; values: [-98.91,-98.83, -98.57]AIUPred-score: 0.56
H1 (60, 1.00e-03); indices(add 1 for resid): 26 85; values: [-98.95,-98.85, -97.57]AIUPred-score: 0.37
H1 (78, 1.00e-03); indices(add 1 for resid): 220 297; values: [-98.94,-98.80, -96.17]AIUPred-score: 0.38
* H1 (22, 5.00e-03); indices(add 1 for resid): 3 24; values: [-98.92,-98.43, -96.03]AIUPred-score: 0.94
H1 (161, 1.00e-03); indices(add 1 for resid): 137 297; values: [-98.96,-98.81, -96.10]AIUPred-score: 0.16
H1 (316, 1.00e-03); indices(add 1 for resid): 0 315; values: [-98.96,-98.79, -84.52]AIUPred-score: 0.10
Processing group High with H0:0 and H1:0 cases
Processing group Low with H0:0 and H1:0 cases
Processing group Very low with H0:0 and H1:0 cases
```

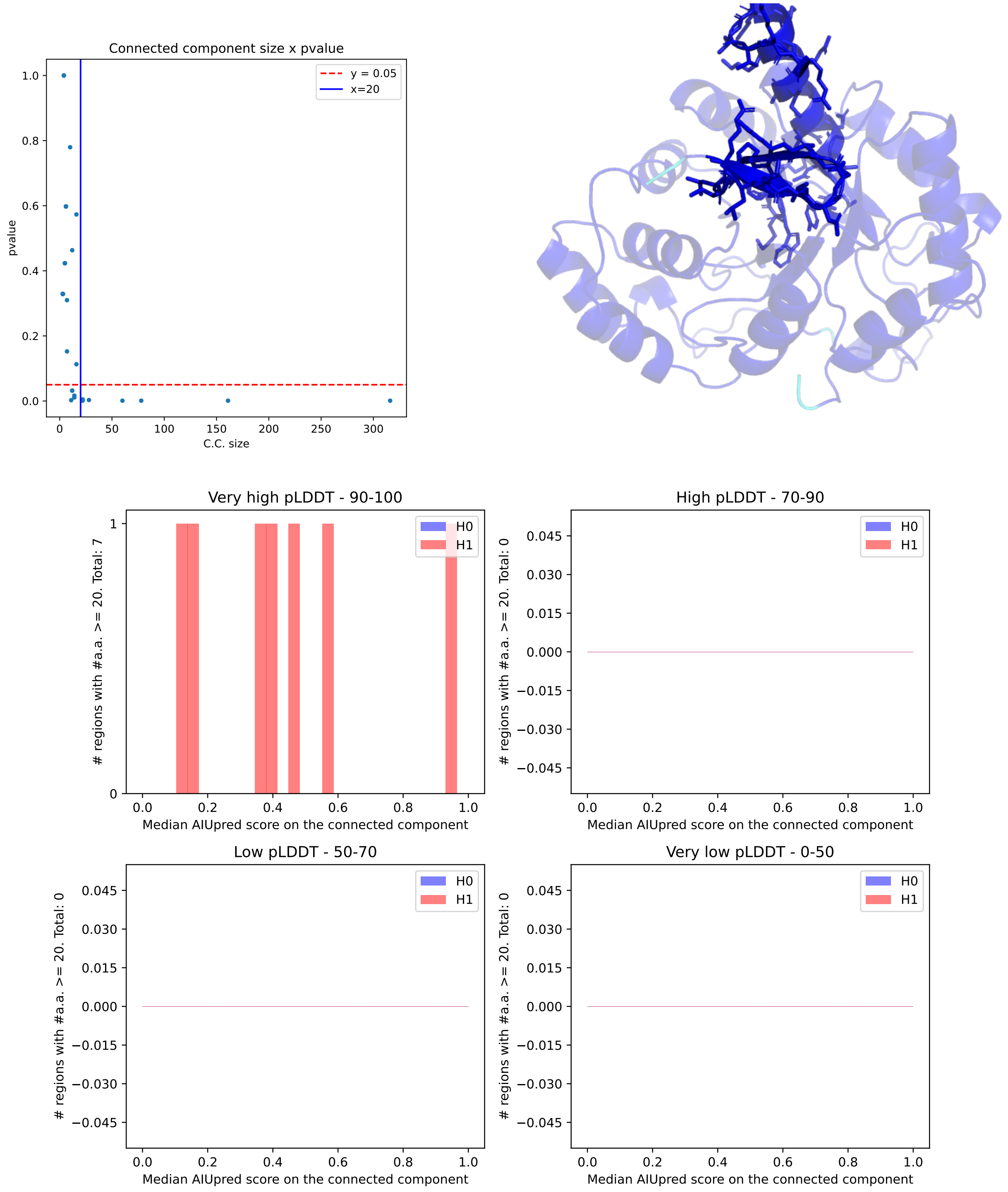

Figure S8: **Model AF-P15121-F1-model-v4. Polyptych plot with minimum connected component size of 20. (First row)** Scatter plot for pvalues and structure. Regions with sticks correspond to c.c. marked with a * in Table S2 **(Second row)** Polyptych plot.

**Table of p-values for connected components S3** Input file AF-Q02817-F15-model_v4.pdb

```
sbl-alphafold-dbrun.py --pval-cc-size 20 --aiu  -f human-AF-Q02817-F15-model_v4.pdb
...
Processing group Very high with H0:0 and H1:0 cases
Processing group High with H0:0 and H1:0 cases
Processing group Low with H0:0 and H1:0 cases
Processing group Very low with H0:26 and H1:25 cases
H0 (39, 8.55e-01); indices(add 1 for resid): 619 657; values: [-40.93,-37.63, -33.65] AIUPred-score: 0.97
H0 (23, 7.70e-01); indices(add 1 for resid): 589 611; values: [-39.31,-35.95, -32.43] AIUPred-score: 0.91
H0 (22, 1.22e-01); indices(add 1 for resid): 1072 1093; values: [-42.77,-38.23, -32.53] AIUPred-score: 0.89
H0 (21, 1.68e-01); indices(add 1 for resid): 526 546; values: [-37.44,-34.55, -28.25] AIUPred-score: 0.87
H0 (22, 5.11e-01); indices(add 1 for resid): 1095 1116; values: [-40.46,-35.32, -31.19] AIUPred-score: 0.89
H0 (223, 3.50e-01); indices(add 1 for resid): 526 748; values: [-41.80,-36.11, -28.25] AIUPred-score: 0.94
H0 (20, 9.67e-01); indices(add 1 for resid): 1118 1137; values: [-35.58,-32.68, -27.97] AIUPred-score: 0.88
H0 (23, 4.84e-01); indices(add 1 for resid): 457 479; values: [-34.59,-32.05, -27.22] AIUPred-score: 0.93
H0 (25, 5.86e-01); indices(add 1 for resid): 114 138; values: [-45.49,-37.58, -27.01] AIUPred-score: 0.94
H0 (44, 3.31e-01); indices(add 1 for resid): 457 500; values: [-35.51,-32.11, -26.39] AIUPred-score: 0.97
H0 (22, 4.24e-01); indices(add 1 for resid): 313 334; values: [-40.20,-33.20, -27.07] AIUPred-score: 0.89
H0 (63, 5.30e-01); indices(add 1 for resid): 457 519; values: [-36.07,-32.31, -26.10] AIUPred-score: 0.98
H0 (21, 1.68e-01); indices(add 1 for resid): 434 454; values: [-41.08,-32.66, -26.07] AIUPred-score: 0.87
H0 (56, 1.72e-01); indices(add 1 for resid): 83 138; values: [-50.56,-41.03, -26.52] AIUPred-score: 0.98
H0 (23, 8.88e-01); indices(add 1 for resid): 234 256; values: [-37.68,-30.40, -25.80] AIUPred-score: 0.90
H0 (23, 1.43e-01); indices(add 1 for resid): 184 206; values: [-38.28,-34.65, -25.69] AIUPred-score: 0.93
H0 (22, 1.95e-01); indices(add 1 for resid): 382 403; values: [-40.61,-32.83, -25.68] AIUPred-score: 0.89
H0 (26, 6.65e-02); indices(add 1 for resid): 405 430; values: [-37.92,-32.20, -25.58] AIUPred-score: 0.94
H0 (20, 7.51e-01); indices(add 1 for resid): 773 792; values: [-37.35,-29.23, -25.98] AIUPred-score: 0.88
H0 (37, 3.35e-01); indices(add 1 for resid): 234 270; values: [-40.09,-30.80, -24.41] AIUPred-score: 0.97
H0 (23, 3.77e-01); indices(add 1 for resid): 819 841; values: [-35.98,-28.07, -24.29] AIUPred-score: 0.91
H0 (22, 4.24e-01); indices(add 1 for resid): 161 182; values: [-40.07,-34.09, -24.45] AIUPred-score: 0.92
H0 (23, 2.74e-01); indices(add 1 for resid): 434 456; values: [-42.32,-32.66, -24.03] AIUPred-score: 0.93
H0 (33, 4.60e-01); indices(add 1 for resid): 1227 1259; values: [-41.65,-29.84, -24.01] AIUPred-score: 0.96
H0 (43, 2.84e-01); indices(add 1 for resid): 140 182; values: [-44.71,-31.78, -23.91] AIUPred-score: 0.97
H0 (23, 4.84e-01); indices(add 1 for resid): 796 818; values: [-34.40,-27.38, -22.48] AIUPred-score: 0.91
H1 (22, 9.00e-03); indices(add 1 for resid): 1049 1070; values: [-47.60,-40.77, -34.89]AIUPred-score: 0.89
H1 (23, 3.50e-03); indices(add 1 for resid): 1003 1025; values: [-47.46,-39.76, -32.49]AIUPred-score: 0.91
H1 (23, 1.00e-03); indices(add 1 for resid): 980 1002; values: [-45.81,-38.36, -31.82]AIUPred-score: 0.91
H1 (23, 3.75e-02); indices(add 1 for resid): 566 588; values: [-38.86,-35.24, -30.67]AIUPred-score: 0.91
H1 (23, 1.00e-03); indices(add 1 for resid): 957 979; values: [-42.88,-36.12, -29.93]AIUPred-score: 0.91
H1 (23, 3.50e-03); indices(add 1 for resid): 934 956; values: [-39.84,-34.15, -28.84]AIUPred-score: 0.91
H1 (23, 3.35e-02); indices(add 1 for resid): 83 105; values: [-49.97,-41.87, -28.79]AIUPred-score: 0.91
H1 (23, 3.50e-03); indices(add 1 for resid): 911 933; values: [-37.28,-32.79, -28.09]AIUPred-score: 0.91
H1 (23, 3.50e-03); indices(add 1 for resid): 865 887; values: [-33.58,-29.73, -25.56]AIUPred-score: 0.91
H1 (89, 4.00e-03); indices(add 1 for resid): 50 138; values: [-51.78,-39.29, -25.87]AIUPred-score: 0.98
H1 (26, 8.00e-03); indices(add 1 for resid): 1325 1350; values: [-40.21,-32.78, -25.81]AIUPred-score: 0.94
H1 (72, 1.50e-03); indices(add 1 for resid): 359 430; values: [-43.45,-32.48, -25.21]AIUPred-score: 0.98
H1 (23, 3.35e-02); indices(add 1 for resid): 1279 1301; values: [-40.86,-31.57, -24.67]AIUPred-score: 0.91
H1 (23, 3.50e-03); indices(add 1 for resid): 842 864; values: [-34.70,-29.26, -24.57]AIUPred-score: 0.91
H1 (72, 2.00e-03); indices(add 1 for resid): 1279 1350; values: [-41.92,-32.35, -24.67]AIUPred-score: 0.98
H1 (92, 2.30e-02); indices(add 1 for resid): 1260 1351; values: [-49.99,-32.32, -24.35]AIUPred-score: 0.98
H1 (104, 3.45e-02); indices(add 1 for resid): 234 337; values: [-42.60,-31.49, -24.28]AIUPred-score: 0.98
H1 (50, 5.00e-03); indices(add 1 for resid): 184 233; values: [-41.83,-32.45, -24.12]AIUPred-score: 0.98
H1 (30, 1.00e-02); indices(add 1 for resid): 1139 1168; values: [-37.74,-29.88, -24.61]AIUPred-score: 0.96
H1 (361, 1.00e-03); indices(add 1 for resid): 434 794; values: [-42.92,-34.23, -24.03]AIUPred-score: 0.92
H1 (414, 1.00e-03); indices(add 1 for resid): 796 1209; values: [-48.73,-32.95, -22.48]AIUPred-score: 0.92
H1 (612, 1.00e-03); indices(add 1 for resid): 184 795; values: [-47.08,-33.59, -21.56]AIUPred-score: 0.92
H1 (144, 1.00e-03); indices(add 1 for resid): 40 183; values: [-51.93,-35.56, -21.10]AIUPred-score: 0.96
H1 (40, 9.00e-03); indices(add 1 for resid): 0 39; values: [-53.05,-41.91, -16.16]AIUPred-score: 0.97
H1 (1400, 1.00e-03); indices(add 1 for resid): 0 1399; values: [-55.36,-33.54, -16.16]AIUPred-score: 0.92
```

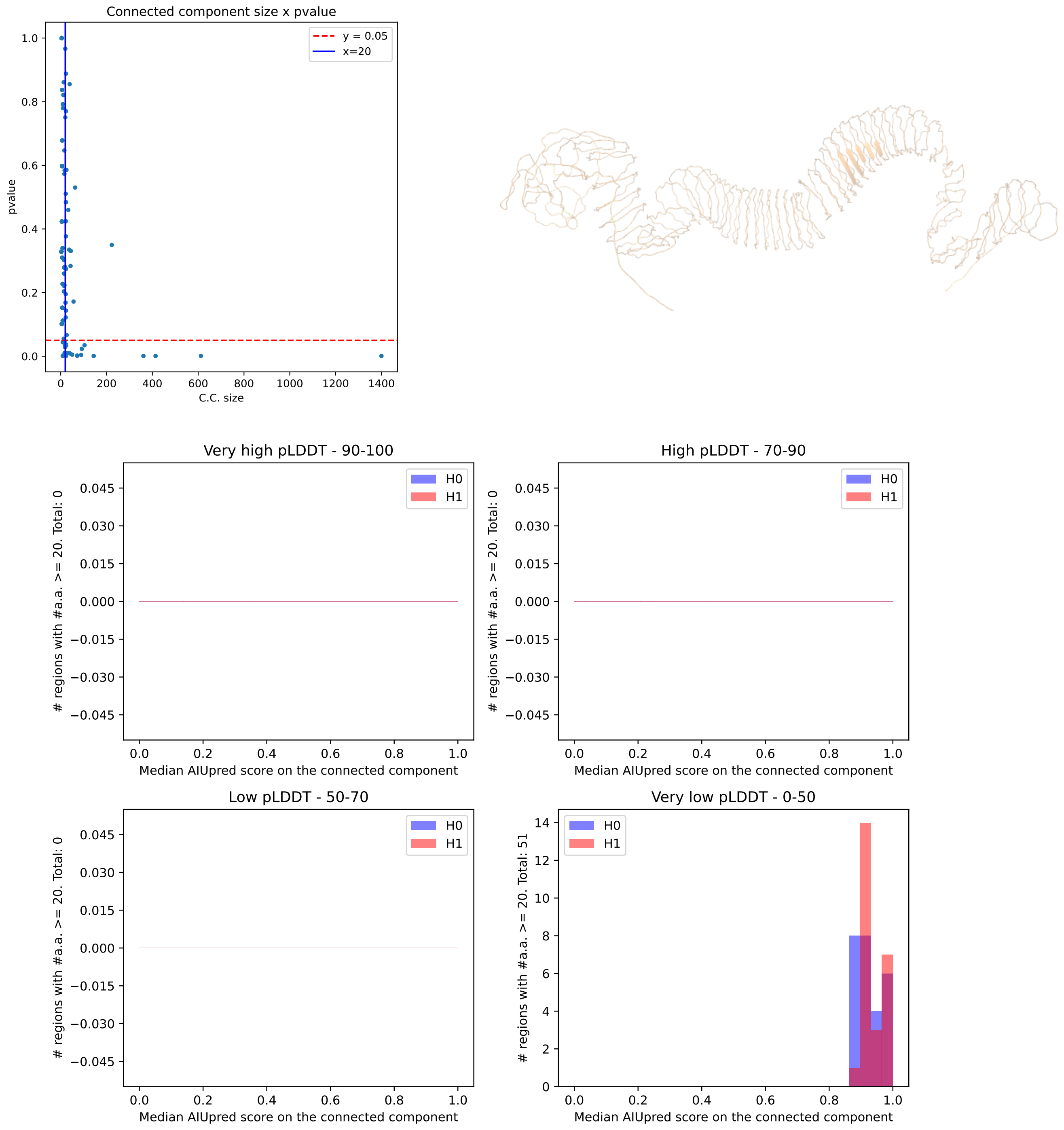

Figure S9: **Model AF-Q02817-F15-model-v4. Polyptych plot with minimum connected component size of 20. (First row)** Scatter plot for pvalues and structure. NB: this example does not display incoherent c.c. (high pLDDT+high `AIUPred` scores or (low pLDDT+low `AIUPred` scores) **(Second row)** Polyptych plot.

**Table of p-values for connected components S4** (Drosophilia) Input file AF-Q9VQS4-F1-model_v4.pdb

```
sbl-alphafold-dbrun.py --pval-cc-size 20 --aiu  -f droso-AF-Q9VQS4-F1-model_v4.pdb
...
Processing group Very high with H0:1 and H1:1 cases
* H0 (41, 3.81e-01); indices(add 1 for resid): 178 218; values: [-95.43,-92.19, -87.13] AIUPred-score: 0.61
* H1 (92, 1.00e-03); indices(add 1 for resid): 130 221; values: [-95.63,-91.98, -73.02]AIUPred-score: 0.61
Processing group High with H0:0 and H1:4 cases
* H1 (53, 1.00e-03); indices(add 1 for resid): 277 329; values: [-93.55,-88.22, -79.07]AIUPred-score: 0.53
H1 (75, 1.00e-03); indices(add 1 for resid): 336 410; values: [-92.67,-87.04, -62.59]AIUPred-score: 0.47
H1 (199, 1.00e-03); indices(add 1 for resid): 231 429; values: [-96.04,-86.80, -49.58]AIUPred-score: 0.44
* H1 (71, 1.00e-03); indices(add 1 for resid): 0 70; values: [-95.39,-87.99, -37.30]AIUPred-score: 0.57
Processing group Low with H0:0 and H1:1 cases
H1 (781, 1.00e-03); indices(add 1 for resid): 0 780; values: [-96.25,-60.74, -34.60]AIUPred-score: 0.58
Processing group Very low with H0:9 and H1:1 cases
H0 (33, 9.17e-01); indices(add 1 for resid): 608 640; values: [-55.59,-47.33, -41.34] AIUPred-score: 0.95
H0 (53, 4.52e-01); indices(add 1 for resid): 592 644; values: [-56.46,-48.10, -38.67] AIUPred-score: 0.97
H0 (24, 2.73e-01); indices(add 1 for resid): 439 462; values: [-50.68,-42.88, -39.10] AIUPred-score: 0.91
H0 (21, 1.68e-01); indices(add 1 for resid): 558 578; values: [-52.49,-45.84, -37.48] AIUPred-score: 0.81
H0 (68, 2.07e-01); indices(add 1 for resid): 660 727; values: [-56.78,-45.02, -37.62] AIUPred-score: 0.87
H0 (171, 8.33e-01); indices(add 1 for resid): 558 728; values: [-57.56,-46.17, -37.13] AIUPred-score: 0.97
H0 (39, 1.47e-01); indices(add 1 for resid): 464 502; values: [-54.79,-43.14, -36.55] AIUPred-score: 0.77
H0 (33, 8.77e-01); indices(add 1 for resid): 504 536; values: [-53.08,-43.61, -36.54] AIUPred-score: 0.69
H0 (234, 2.00e-01); indices(add 1 for resid): 547 780; values: [-59.92,-46.05, -36.27] AIUPred-score: 0.98
H1 (25, 1.15e-02); indices(add 1 for resid): 666 690; values: [-55.45,-45.27, -38.59]AIUPred-score: 0.96
```

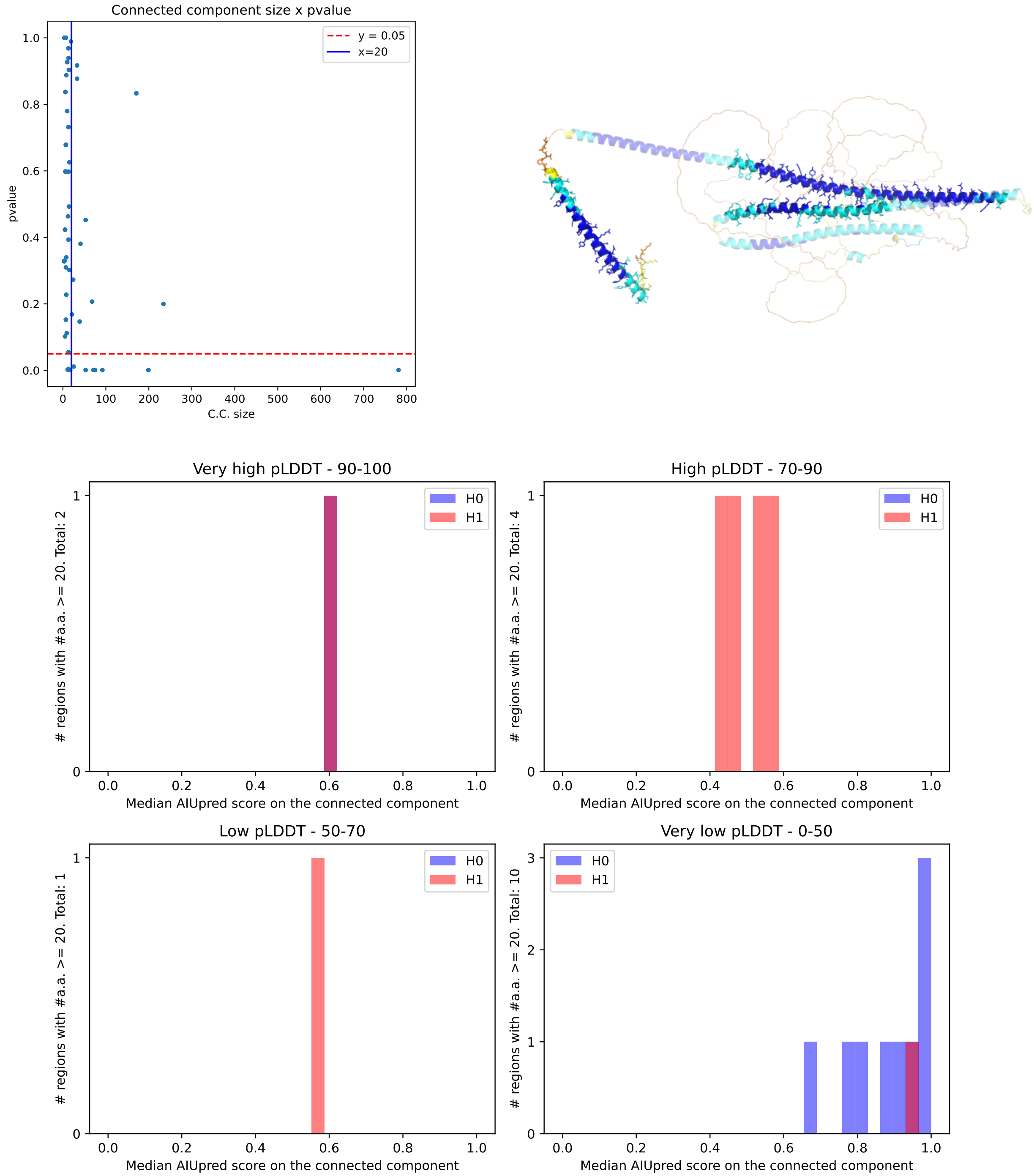

Figure S10: **Model AF-Q9VQS4-F1-model-v4. Polyptych plot with minimum connected component size of 20. (First row)** Scatter plot for pvalues and structure. Regions with sticks correspond to c.c. marked with a * in Table S4 **(Second row)** Polyptych plot.

**Table of p-values for connected components S5** Input file AF-Q9VQS4-F1-model_v4.pdb

```
sbl-alphafold-dbrun.py --aiu -f AF-A0A0H2YV13-F1-model_v6.pdb
...
Processing group Very high with H0:0 and H1:5 cases
H1 (25, 1.00e-03); indices(add 1 for resid): 1011 1035; values: [-98.12,-94.38, -89.12]AIUPred-score: 0.40
H1 (29, 2.50e-03); indices(add 1 for resid): 803 831; values: [-96.88,-93.31, -84.44]AIUPred-score: 0.51
H1 (21, 1.50e-03); indices(add 1 for resid): 944 964; values: [-96.19,-91.81, -84.50]AIUPred-score: 0.85
* H1 (74, 1.00e-03); indices(add 1 for resid): 803 876; values: [-98.38,-93.69, -79.94]AIUPred-score: 0.63
* H1 (56, 1.00e-03); indices(add 1 for resid): 991 1046; values: [-98.69,-94.00, -79.69]AIUPred-score: 0.34
Processing group High with H0:1 and H1:13 cases
H0 (23, 4.84e-01); indices(add 1 for resid): 967 989; values: [-95.50,-88.94, -80.75] AIUPred-score: 0.47
* H1 (21, 3.15e-02); indices(add 1 for resid): 647 667; values: [-86.69,-71.62, -60.16]AIUPred-score: 0.88
H1 (26, 8.00e-03); indices(add 1 for resid): 216 241; values: [-88.38,-81.50, -55.94]AIUPred-score: 0.45
* H1 (30, 1.00e-02); indices(add 1 for resid): 45 74; values: [-87.12,-74.31, -51.84]AIUPred-score: 0.91
H1 (42, 1.50e-03); indices(add 1 for resid): 215 256; values: [-88.94,-81.72, -50.75]AIUPred-score: 0.34
* H1 (31, 1.00e-03); indices(add 1 for resid): 718 748; values: [-91.44,-77.44, -48.34]AIUPred-score: 0.75
H1 (24, 1.00e-03); indices(add 1 for resid): 153 176; values: [-86.19,-72.81, -49.06]AIUPred-score: 0.48
H1 (67, 1.00e-03); indices(add 1 for resid): 717 783; values: [-92.88,-79.75, -36.66]AIUPred-score: 0.28
* H1 (66, 1.00e-03); indices(add 1 for resid): 112 177; values: [-89.62,-71.44, -37.34]AIUPred-score: 0.52
H1 (80, 1.00e-03); indices(add 1 for resid): 185 264; values: [-89.38,-76.53, -25.62]AIUPred-score: 0.22
H1 (80, 1.00e-03); indices(add 1 for resid): 630 709; values: [-90.94,-73.59, -22.83]AIUPred-score: 0.44
H1 (271, 1.00e-03); indices(add 1 for resid): 0 270; values: [-89.69,-70.25, -22.81]AIUPred-score: 0.10
H1 (81, 1.00e-03); indices(add 1 for resid): 545 625; values: [-88.69,-70.75, -21.61]AIUPred-score: 0.49
H1 (1049, 1.00e-03); indices(add 1 for resid): 0 1048; values: [-98.75,-75.19, -21.33]AIUPred-score: 0.09
Processing group Low with H0:0 and H1:3 cases
H1 (82, 1.00e-03); indices(add 1 for resid): 375 456; values: [-87.94,-67.16, -23.78]AIUPred-score: 0.50
H1 (84, 1.00e-03); indices(add 1 for resid): 458 541; values: [-88.56,-68.19, -21.94]AIUPred-score: 0.54
H1 (543, 1.00e-03); indices(add 1 for resid): 0 542; values: [-92.19,-69.12, -21.33]AIUPred-score: 0.11
Processing group Very low with H0:1 and H1:0 cases
H0 (20, 1.56e-01); indices(add 1 for resid): 271 290; values: [-49.53,-25.48, -23.05] AIUPred-score: 0.73
```

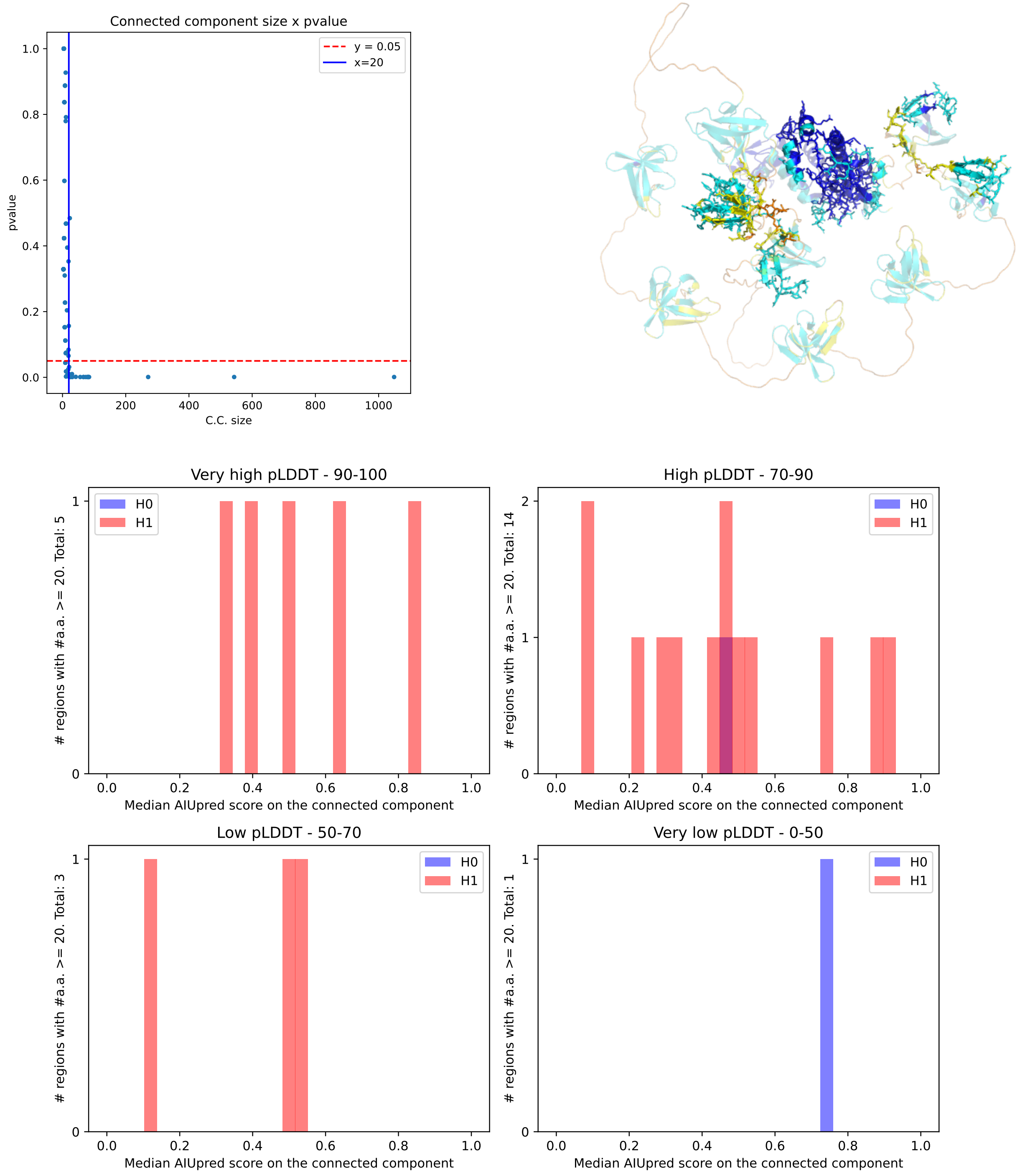

Figure S11: **Model AF-A0A0H2YV13-F1-model. Polyptych plot with minimum connected component size of 20. (First row)** Scatter plot for pvalues and structure. Regions with sticks correspond to c.c. marked with a * in Table S5 **(Second row)** Polyptych plot.

**3 Results** **5**

**4 Outlook** **7**

**5 Artwork** **8**

**S1 SI** **16**
S1.1 pLDDT statistics on whole genomes . . . 16
S1.2 Algorithms . . . 16
S1.3 Proof of theorem 1 . . . 16
S1.4 Results–general . . . 18
